# An ancient retrotransposon provides species-specific tuning of IL-18 inflammatory signaling

**DOI:** 10.64898/2026.05.25.727421

**Authors:** Andrea D. Ordonez, Holly Allen, Lynn Sanford, Atma Ivancevic, Ashley Agyepong, Mia Chaw, James P. Bridges, Tony Schountz, Edward B. Chuong

## Abstract

Species differ markedly in how they regulate immune responses, yet the molecular basis of this variation remains incompletely understood. Here, we characterize an underexplored regulatory mechanism of the pro-inflammatory IL-18 pathway, centered on a truncated *IL18R1* isoform with striking species-specific expression differences. This isoform, *IL18R1-Short*, derives from an ancient LINE2 retrotransposon insertion that provides an intronic polyadenylation signal, producing a receptor that lacks the intracellular signaling domain. Using RNA sequencing across nine mammals, we find that although this cis-regulatory element is broadly conserved, robust *IL18R1-Short* expression is species- and tissue-restricted: bats and mice express it at high levels, particularly in barrier tissues (lung, skin, intestine), whereas most other species, including humans, show little or no expression. Functionally, IL18R1-Short dampens IL-18–induced NFκB signaling in human, mouse, and bat systems, and knockdown of the mouse ortholog enhances IL-18–driven immune and inflammatory gene expression in mouse T cells. Together, these results identify IL18R1-Short as a transposable element-derived decoy receptor and highlight alternative transcription as a source of species-specific immune regulation.

## Introduction

Regulation of immune responses is essential for mounting an effective host defense while minimizing the risk of autoimmunity, chronic inflammation, and cancer ^1–3^. Although the core architecture of immune signaling is highly conserved across mammals, species differ markedly in how they regulate immune responses, leading to striking variation in susceptibility and tolerance to infection or disease ^4–8^. Identifying these species-specific differences in immune regulation is critical for both improving the translational relevance of animal models ^9–11^ and for predicting host responses to emerging pathogens ^12,13^.

Historically, comparative studies of immune evolution have focused on changes to protein-coding sequences, including gene copy number variation and signatures of positive selection within coding domains. For example, analyses of bat genomes have uncovered multiple lineage-specific adaptations including the loss of the PYHIN locus that contains important innate immune sensors ^14^, the contraction of the type I interferon (IFN) locus ^15^, and dampened function mutations in foreign DNA sensor STING ^16^. Moreover, a large body of work has delineated the immune differences between humans and mice including a different repertoire of Natural Killer (NK) cell receptors ^17^, functional toll-like receptors (TLRs) ^17,18^, and cytokines, such as interleukin-8 (IL8) ^19^ and IL37 ^19,20^. More recently, regulatory differences affecting gene expression levels and inducibility have emerged as another major axis of immune diversification ^15,21–26^. Together, these studies highlight multiple routes to immune divergence, but do not address how transcript-level changes in gene architecture—rather than coding mutations or expression level—can shape species-specific immune regulation.

Alternative isoform expression can produce protein variants that lack key functional domains, exhibit altered subcellular localization, or interfere with canonical signaling complexes, thereby fine-tuning immune responses without altering core pathway architecture ^27–29^. These events have been extensively studied in humans where alternative isoforms have been shown to regulate IFN responses ^30,31^, TLR signaling cascades ^32–34^, and cytokine signaling pathways ^35–38^. Comparable work in non-human primates and mice exists but is more limited ^39–43^. As a result, we still lack a clear view of which immune isoforms are conserved, as opposed to lineage- or species-specific, and when isoform usage produces meaningful functional divergence across mammals.

Here, we investigate how alternative transcription contributes to species-specific evolution of immune regulation, focusing on IL-18 signaling. IL-18 is produced by macrophages and barrier epithelial cells as an inactive pro-cytokine that requires inflammasome-dependent caspase cleavage for activation ^44–48^. Canonical IL-18 signaling occurs through a heterodimeric receptor consisting of IL-18 receptor 1 (IL18R1; IL18R1α) and the accessory protein IL18RAP (IL18R1β) ^49^. Ligand binding triggers recruitment of the adaptor MYD88, leading to downstream activation of transcription factors including nuclear factor κB (NFκB) and activator protein 1 (AP-1) ^44,50,51^.

IL-18 is best known for promoting IFNγ-mediated inflammatory responses, particularly by synergizing with IL-12 to enhance NK and T cell activity ^44,51–53^. Loss of proper regulation is linked to hyperinflammatory pathology ^54–56^, with elevated IL-18 secretion and activity reported in macrophage activation syndrome ^57,58^ and NLRC4-driven autoinflammation ^59^. Accordingly, IL-18 signaling is normally modulated by several inhibitory mechanisms. The soluble IL-18 binding protein (IL18BP) neutralizes the ligand and reduces downstream outputs including IFNγ responses in vitro and in vivo ^60,61^, and loss of IL18BP exacerbates inflammatory states in mouse ^62^, including in the context of macrophage activation syndrome ^63^. Regulation also occurs through inhibitory receptor complexes involving SIGIRR ^64^, and loss of this regulator leads to increased intestinal inflammation and enhanced colitis-associated tumorigenesis in mice ^65,66^. In addition, SIGIRR can pair with IL18R1 to bind IL37 and actively induce anti-inflammatory signaling, rather than merely suppressing pro-inflammatory responses ^20,67^. The evolution of multiple regulatory layers establishes the importance of precisely tuning IL-18 signaling to achieve optimal proinflammatory activity.

To date, most research on IL-18 regulation has focused on the proteins described above. However, previous studies implicate alternative transcription at the *IL18R1* locus as an additional mode of regulation, generating a truncated isoform (hereafter *IL18R1-Short*) that negatively regulates this pathway ^68,69^. This isoform is generated by an early polyadenylation signal (PAS) within intron 9, and is predicted to encode a truncated receptor that retains the extracellular ligand-binding domain but lacks the intracellular signaling domain. Expression of *IL18R1-Short* has been detected in the rat lung ^69^ in the mouse brain ^68^, and in a subset of human immune cell types, including plasmacytoid dendritic cells (pDCs) and NK cells ^70^. Additionally, overexpression of human IL18R1-Short was shown to dampen NFκB activity in a reporter cell system, supporting a model in which this isoform functions as a decoy receptor ^70^. Notably, the intronic polyadenylation signal giving rise to IL18R1-Short is preserved across many vertebrate genomes, suggesting an evolutionarily conserved regulatory strategy ^69^.

Despite its potential importance in regulating IL-18 signaling, the IL18R1-Short isoform has received limited attention, with no mechanistic follow-up studies over the past decade. Prior work relied largely on non-quantitative approaches, such as RT-PCR, to detect the isoform. Moreover, because intronic polyadenylation (IPA) events are often suppressed ^71^ or subject to nuclear surveillance and decay ^72^, it has remained unclear whether IL18R1-Short is produced at physiologically meaningful levels. As such, virtually all studies to date on IL-18 signaling implicitly assume expression of only the full-length canonical IL18R1 protein.

Here, we investigate the evolution, expression, and function of IL18R1-Short. We trace the intronic PAS regulating this isoform to an ancient LINE2 (L2) retrotransposon insertion conserved across eutherian and marsupial mammals. Using isoform-resolved transcriptomics across nine species and a broad panel of primary tissues, we show that *IL18R1-Short* is highly expressed in bats and mice but largely absent in most other mammals, including humans. This species-restricted pattern occurs despite broad conservation of the underlying cis element, indicating that the presence of transposable element (TE)-derived regulation is necessary but not sufficient for functional co-option. We demonstrate that IL18R1-Short dampens IL18-induced NFκB signaling in human and mouse systems, and silencing the endogenous *Il18r1-short* isoform in mouse T cells amplifies inflammatory gene expression. Together, these results establish IL18R1-Short as a TE-derived decoy receptor that modulates IL-18 activity in a species- and tissue-specific manner, revealing how latent regulatory elements introduced by ancient transposable element insertions can be differentially co-opted to shape immune signaling across mammalian lineages.

## Results

### An ancient L2-derived intronic PAS enables *IL18R1-Short* expression in a species- and tissue-restricted manner

*IL18R1-Short* is an alternative isoform of the canonical full-length gene *IL18R1*, generated by intronic polyadenylation that truncates the transcript, deleting the coding sequence (CDS) for the intracellular signaling domain (Figure 1A). This isoform has been proposed to be expressed across mammals based on conservation of the predicted PAS motif ^69^. We find that the PAS is provided by an ancient L2 retrotransposon (Figure 1A) conserved across eutherian and marsupial mammals but absent from platypus, consistent with insertion ∼160-186 million years ago (Figure 1B). Alignment of the L2-derived PAS region showed an intact canonical PAS motif (AATAAA) in all species examined except cow (*Bos taurus*) (Figure 1A).

**Figure 1.**
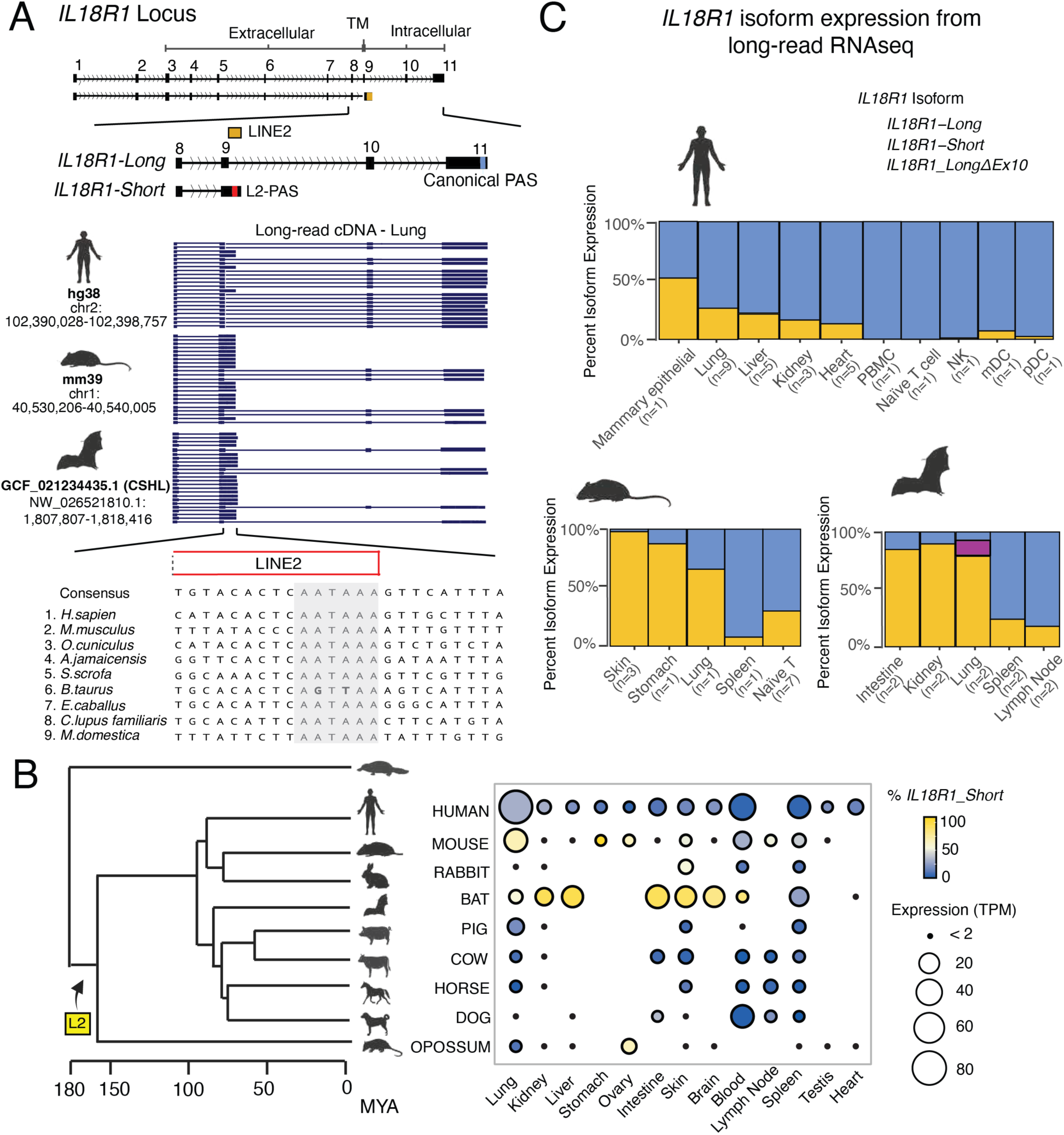
*IL1BR1-Short* is differentially expressed across mammals. **A** Schematic of the *IL18R1* locus showing the canonical full-length isoform *(IL 18R1-Long)* and the truncated isoform *(IL18R1-Short),* which terminates within intron 9 at an intronic PAS contributed by an L2 element. Long-read cDNA alignments from lung are shown for human, mouse, and Jamaican fruit bat; coordinates and genome versions for each locus are given. (Bottom) sequence alignment of the L2-derived PAS region across representative mammals showing conservation of the retrotransposon and associated PAS sequence, highlighted in grey. **B** Phylogenetic tree of mammalian species generated by TimeTree (Kumar et al. 2022), shows predicted time of L2 insertion, with platypus as an outgroup. Average expression of *IL18R1-Short* in percentage of reads quantified for locus from short-read bulk RNA-seq across tissues. Samples with ≤ 2 Transcripts Per Million (TPM) total *IL1BR1* expression are represented as black dots; blank spaces indicate no expression data analyzed for that tissue; color of bubble represents isoform expression: yellow - high *IL18R1-Short* and low *IL 18R1-Long* and blue - high *IL 18R1-Long* and low *IL18R1-Short.* CAverage percent expression of long and short *IL18R1* isoforms across tissues from long-read RNA-seq. Blue= *IL18R1-Long,* yellow= *IL18R1-Short* and purple= *IL18R1_LonglJ.Ex10.* Percent expression of *IL18R1-Shortor IL18R1-Long* was calcu-

Previous studies detected *IL18R1-Short* in a limited set of species and tissues (mouse brain, rat lung, and human NK cells and pDCs) using low-throughput approaches, including RT-PCR, Sanger sequencing, and NanoString nCounter expression profiling ^69,70^. However, whether expression of the short isoform is broadly conserved across mammals—and whether it is physiologically relevant—remains unclear. To address this, we analyzed *IL18R1* isoform-level expression using bulk RNA sequencing (RNA-seq) across a panel of tissues from nine representative mammals. This analysis integrated publicly available datasets with in-house generated data for the Jamaican fruit bat (*Artibeus jamaicensis*) (Table S1 and 3A). Across species, *IL18R1* locus-level expression was moderate to robust in various tissues (Figure 1B). As expected *IL18R1-Long* showed high expression in immune tissues, including blood, lymph node, and spleen (Figure 1B; Figure S1). In comparison, *IL18R1-Short* was largely absent (Figure 1B), generally comprising 0–20% of total locus expression (Table S2). In human, our most sampled species, *IL18R1-Short* was widely undetectable in all tissues except for lung (28.64%; Figure 1B; Table S2). These results indicate that *IL18R1-Short* is unlikely to be functional in most mammals, given its generally low expression in the tissues examined, consistent with the tendency of IPA-derived transcripts to be repressed.

Notably, we detected high *IL18R1-Short* expression (>50% of total locus expression) in a restricted subset of species—Jamaican fruit bat, mouse, rabbit, and opossum (Figure 1B; Figure S1). In both rabbit and opossum, we only detected high *IL18R1-Short* expression in one tissue; rabbit skin (50.27%) and opossum ovary (60.03%) (Figure 1B; Figure S1). Surprisingly, bat *IL18R1-Short* was highly expressed in multiple tissues, including blood (78.9%), intestine (86.7%), liver (89.5%), kidney (87.4%), skin (84.7%), and brain (74.4%), and more moderately expressed in the lung (51%) (Figure 1B; Table S2). Similarly, mouse *IL18R1-Short* was highly expressed in stomach (96%), lung (61.4%), and ovary (59.4%), showing more moderate expression in skin (47.6%) and lymph node (48.3%) (Figure 1B; Table S2). The unusually strong *IL18R1-Short* expression in Jamaican fruit bat and mouse prompted us to ask whether this pattern extended to other bats and rodents. *IL18R1-Short* was detected in some of these species but showed a patchy expression pattern across tissues overall (Figure S2), likely due to limited sampling.

IPA events can be difficult to quantify accurately by short-read RNA-seq alone ^40,73–75^, therefore we next used long-read RNA-seq to resolve *IL18R1* isoform structure and perform unbiased quantification (Table S3A). We focused on mouse and Jamaican fruit bat, where short-read data indicated robust *IL18R1-Short* expression, and included human as a representative species in which the short isoform is low or absent in most tissues. In each case, long-read sequencing confirmed that *IL18R1-Long* and *IL18R1-Short* are the principal isoforms at this locus. We identified multiple transcripts for each isoform (Figure S3); however, among expressed transcripts (Counts per million (CPM) ≥1; Table S3B), differences between sequences were confined to untranslated regions (UTRs). Notably, Jamaican fruit bat also expressed an additional lung isoform, *IL18R1-LongΔEx10* (Figure 1A, C; Figure S3), which terminates at the canonical PAS but skips exon 10. This alternative splicing event truncates the TIR domain required for MYD88 recruitment and may encode a signaling-deficient receptor.

Our long-read quantification was concordant with short-read RNA-seq and confirmed the species- and tissue-restricted pattern of *IL18R1-Short* expression (Figure 1C; Table S4). In human, *IL18R1-Long* remained the predominant isoform across most tissues, with *IL18R1-Short* comprising 0–25% of transcripts (Figure 1C; Figure S4). The highest *IL18R1-Short* expression was observed in mammary epithelial cells (∼50%) and lung (∼30%) (Figure 1C). In mouse and Jamaican fruit bat, long-read data confirmed enrichment of *IL18R1-Short* in barrier and epithelial tissues (mouse: skin, stomach, and lung; bat: intestine, kidney, and lung; Figure 1C). Beyond barrier tissues, mouse naïve T cells also showed substantial *IL18R1-Short* expression (17–57% across samples), whereas this isoform was not detected in human naïve T cells (Figure 1C; Table S4; Figure S4), suggesting that *IL18R1-Short* regulation may extend to T cell biology in mouse but not in human.

Together, our RNA-seq analysis shows that preservation of the ancient L2-derived PAS is necessary but not sufficient for *IL18R1-Short* expression across mammals and points instead to lineage- and tissue-specific transcriptional and post-transcriptional regulation as key determinants of its expression.

### Cell-type specific expression of *IL18R1-Short* differs between human and mouse lung

The robust expression of *IL18R1-Short* in some species suggests that it may have an adaptive physiological function. To further explore the expression pattern of IL18R1-Short at the cell population level, we first examined the Tabula Muris single-cell dataset ^76^ (SMART-Seq) spanning 20 mouse organs (Figure S5). Immune populations, such as NK and T cells expressed both *Il18r1-long* and *Il18r1-short* at comparable levels (Figure 2A). In contrast, epithelial populations — including lung, mammary gland, epidermis, and large intestine — showed prominent expression of *Il18r1-short* only (Figure 2A), consistent with the barrier-tissue enrichment observed in bulk RNA-seq.

**Figure 2.**
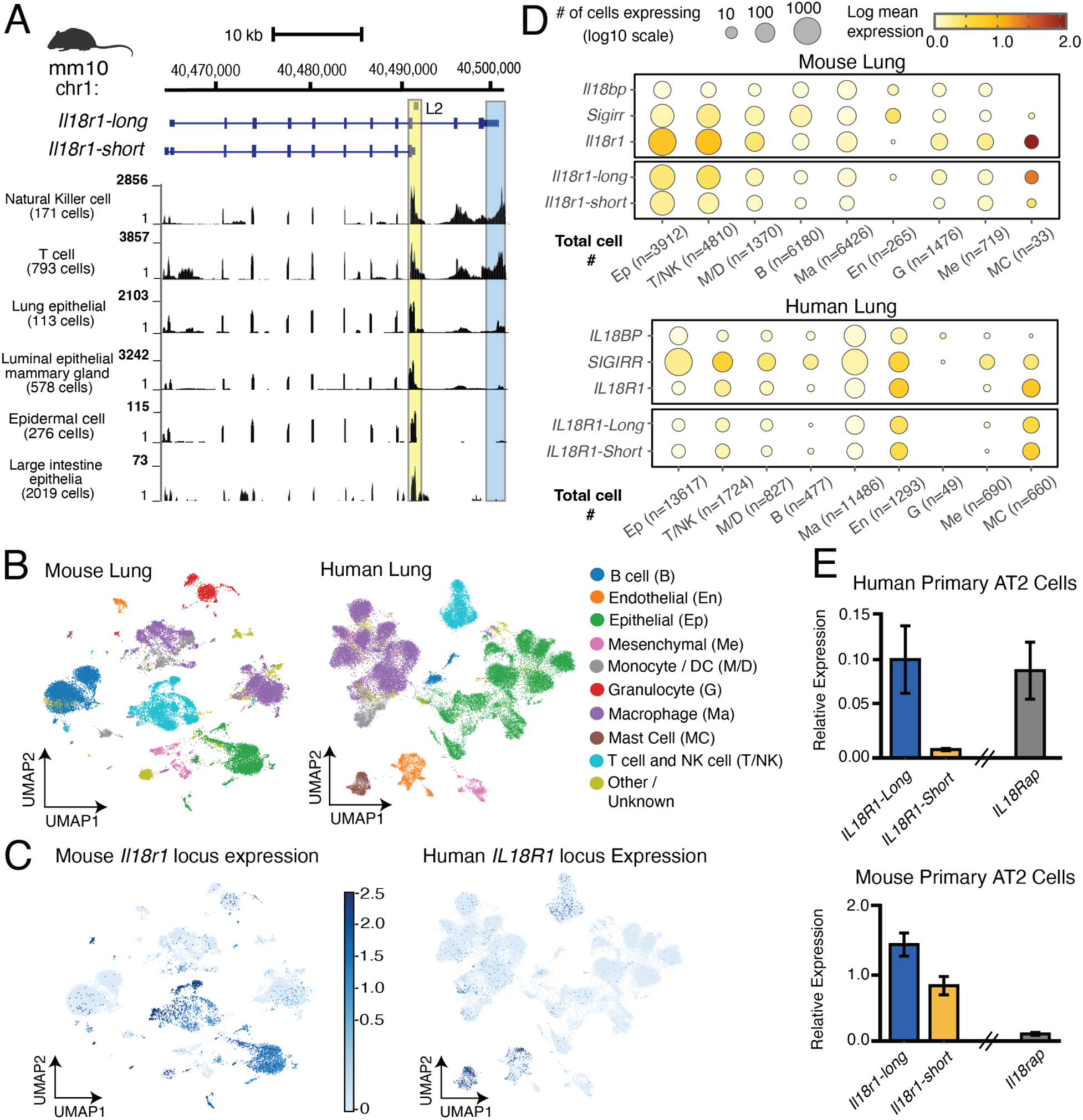
IL 18R1 isoform expression is cell-type specific and enriched in epithelial populations. **A** UCSC Genome Browser tracks (mm10, chr1:40,460,000-40,505,000) showing single-cell RNA-seq coverage for indicated Tabula Muris cell populations (SMART-seq). Il18r1-long and Il18r1-short isoform structures are shown above; the intronic L2-PAS insertion site is highlighted in yellow and the canonical Il18r1-long PAS in blue. Read depth scale (1 to indicated maximum) is shown for each population; cell numbers are indicated in parentheses. **B** Uniform Manifold Approximation and Projection (UMAP) visualization of lung cell populations profiled by 10x Genomics single-cell RNA-seq in mouse (left) and human (right). Cell types are indicated by color; abbreviations are provided for each population. **C** UMAP plots showing total normalized expression of Il18r1 (mouse) and IL18R1 (human) in blue across profiled lung cell populations. Scale bar represents log1p(counts per 10K). UMAPs for Sigirr. Il18bp, and their human orthologs are shown in Supplementary Figs. 6-7. **D** lsoform-resolved expression of Il18r1 (mouse, top) and IL18R1 (human, bottom) across lung cell populations, alongside negative regulators of IL18 signaling (IL18BP and SIGIRR). Circle size reflects the number of cells expressing each gene (log10 scale); circle color indicates log1p mean normalized expression. Cell population labels indicate abbreviated cell type and total cell number (n) for each cluster. **E** lsoform-specific RT-qPCR in primary human (top) and mouse (bottom) AT2 cells from four biological replicates. Relative expression of IL18R1-Long, IL18R1-Short, and IL18RAP was normalized to the housekeeping gene CTCF (2^^^-ΔCq). Bars represent mean ± SEM. Note the different y-axis scales between panels, reflecting higher overall IL18R1 expression in mouse AT2 cells.

We also examined *IL18R1-Short* expression in human lung, where the isoform was detectable by bulk RNA-seq in multiple samples despite being absent from most other tissues (Figure 2B–D; Table S1). Across populations, very few cells expressed the *IL18R1* locus in human (Figure 2C), complicating our quantification of *IL18R1-Short* (Figure 2D). This is likely due to low sampling of this locus, rather than true endogenous expression levels, as bulk RNA-seq of various human lung samples showed high total locus expression for this gene (Figure 1B; Figure S1). Human endothelial and mast cell populations showed the highest percentage of cells expressing *IL18R1*, 12.5% (162 cells) and 16% (104 cells), respectively (Figure 2C–D; Table S8). In both, *IL18R1-Short* and *IL18R1-Long* were expressed at similar levels (Figure 2D; Figure S7), suggesting that the short isoform is more prominent in certain human lung cell types than indicated by bulk RNA-seq. From mouse lung single-cell data, we detected high expression levels of *Il18r1 locus* in epithelial cells (51.6%) and moderate levels in T and NK cells (22.7%) (Figure 2D; Table S7). While the Tabula Muris data showed equivalent expression of short and long *Il18r1* isoforms in T and NK cells and predominant expression of the short isoform in epithelial cells (Figure 2A), the 10X genomics data showed less significant expression of *Il18r1-short* in both cases (Figure 2D; Figure S6). Mast cells in mouse lung showed the highest expression of *Il18r1* (78.8%; Table S7) locus, yet only 33 cells total were captured for this population, and the long isoform is predominant in this cluster (Figure 2D).

To resolve the discrepancies in the expression of *Il18r1-short* in mouse lung epithelial cells from 10X genomics and SMART-Seq data, we performed targeted qPCR on RNA from primary mouse alveolar type 2 (AT2) epithelial cells (Figure 2E, Table S11). *Il18r1-short* was significantly expressed in mouse AT2s, albeit was not the predominant isoform as suggested by Tabula Muris. However, the Tabula Muris cluster specifies lung epithelial (Figure 2A) which encapsulates more than just AT2s. Moreover, the same analysis of primary human AT2s showed much lower normalized expression of *IL18R1* locus compared to the mouse AT2s (Figure 2E; over 10-fold), and the short isoform was barely detectable, matching our observations from the single-cell human lung data (Figure 2D).

Since IL18R1-Short has been hypothesized to function as a regulator of IL-18 ^70^, we compared the expression profiles of *IL18R1-Short* against other known negative regulators of IL-18 in the lung (Figure 2D; Figure S6-7). *SIGIRR* was widely expressed across cell populations in human and mouse lung; in most clusters, it was the most highly expressed negative regulator (Figure 2D; Figure S6-7). In comparison, *IL18BP* is widely but lowly detected across lung cell populations. These different expression profiles suggest context dependent, non-redundant roles for these regulators of IL18. Taking together the single-cell and AT2 qPCR analysis of mouse *Il18r1* isoform expression, *Il18r1-short* likely plays a unique modulatory role in mouse Il-18 signaling at the epithelial cell layer in the lung, and likely other barrier tissues as well.

Lastly, we attempted to analyze a Jamaican fruit bat intestinal single-cell dataset, as no lung single-cell data set exists for this species, and the intestine was a tissue that showed predominant *IL18R1-Short* expression from bulk RNA-seq (Figure 1A). Our analysis only detected significant levels of *IL18R1* in enterocytes, where isoform quantification was challenging due to low coverage of 3’ UTRs. Challenges with cell clustering and annotation in a non-model organism further limited interpretation (Figure S8).

### IL18R1-Short dampens NFκB signaling in human, mouse, and bat IL-18 reporter systems

*IL18R1-Long* encodes canonical IL18R1, a type I transmembrane receptor of IL-18 that requires the accessory receptor IL18RAP for signal transduction. The *IL18R1-Short* isoform is predicted to generate a truncated receptor lacking most of the intracellular signaling domain (Figure 1A) and has been shown to negatively regulate IL-18 NFκB activity when overexpressed in HEK293T cells ^70^. The truncation is conserved across species, differing only in the number of L2-derived amino acids (aa) appended after exon 9 (human: 22 aa; bat: 23 aa; mouse: 5 aa) (Figure S9). We hypothesized that despite these differences in length, *IL18R1-Short* evolved to function as a decoy receptor across species, dampening IL-18 signaling due to the loss of the domain required for MYD88 recruitment.

To test whether *IL18R1-Short* encodes a functional protein, we expressed the human isoform in HEK293T cells and used HaloTag labeling to assess its localization. Fluorescence imaging showed that HaloTag–IL18R1-Short produced a stable protein that localized to the plasma membrane (Figure 3A). In the same system, HaloTag–IL18R1-Long and HaloTag–IL18Rap also localized to the membrane, consistent with their established roles in receptor signaling (Figure 3A; Figure S10).

**Figure 3.**
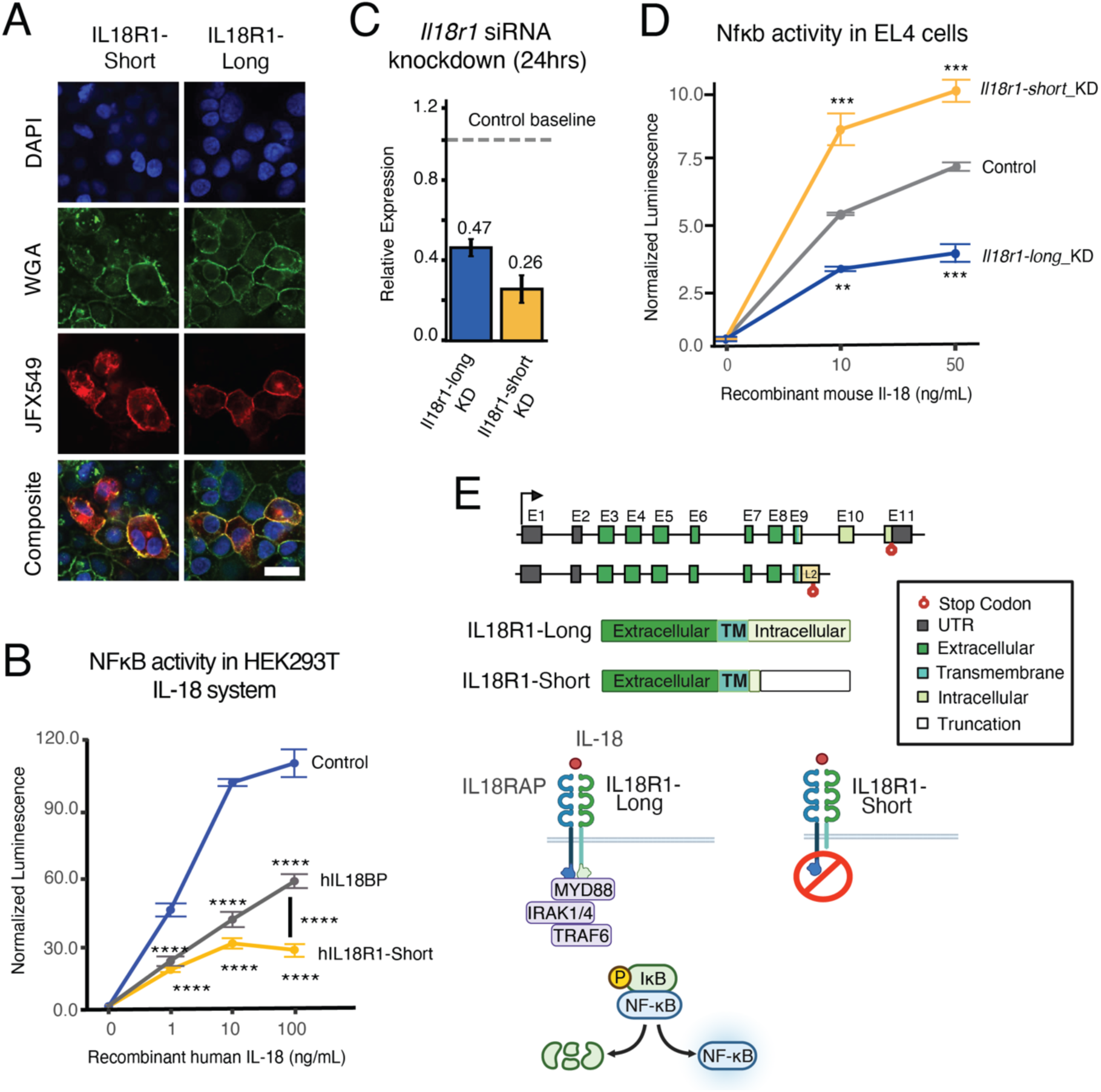
IL18R1·Short is a truncated transmembrane receptor that dampens IL-18-induced NFκB activity. **A** Representative fluorescence images of HEK293T cells transfected with HaloTagged constructs of human IL18R1-Long or IL18R1-Short and stimulated with 100 ng recombinant human IL-18 for 30 min. Images are shown from top to bottom: 405 nm (DAPI), 488 nm (WGA; plasma membrane stain), 561 nm (Janelia Fluor 549; HaloTag ligand), and merged channels. Scale bar, 50 *µm.* **B** IL-18-mediated NFκB activity in human HEK293T cells measured by dual-luciferase reporter assay across increasing concentrations of recombinant IL-18. Control reaction: human IL18R1-Long + IL18RAP; test reactions: human IL18R1-Long + IL18RAP + human IL18R1-Short (500ng) or IL18BP (500ng). NanoLuc luciferase values were normalized to Firefly. Error bars represent ± standard error (SE) of the mean; N = 6 treatment replicates. Significance values shown correspond to test condition vs control, unless otherwise specified. **C** Relative expression of mouse *ll18r1* isoforms 24h after siRNA transfection in EL4 cells, normalized to *Ctcf* and to the non-targeting control (2^ΔΔCq). N = 2 qPCR replicates; KD = siRNA knockdown. **D** Mouse 11-18-mediated Nfκb activity in EL4 cells measured by dual-luciferase reporter assay across increasing concentrations of recombinant Il-18. NanoLuc (NL) luciferase values are normalized to Firefly luciferase (FF); NUFF ratios are normalized to untreated conditions. Error bars represent ± SE of the mean; N= 3 treatment replicates. Significance values shown correspond to isoform KD vs non-targeting control. **E** *IL1BR1* isoform maps and schematic representation of IL18R1-Long and IL18R1-Short receptor proteins and their signaling capacities. Intracellular signaling cascade is depicted culminating in NFκB activation and signaling. P-values were calculated using unpaired two-tailed I-tests(**, p ≤ 0.001;***, p ≤ 0.001;;***, p ≤ 0.0001).

We next used a previously described HEK293T reporter cell system for IL-18–induced NFκB activity, in which IL18R1-Long and IL18RAP are co-expressed to reconstitute human IL-18 responsiveness^70^, and asked whether expression of *IL18R1-Short* alters signaling output. Consistent with the findings of Chao et al. (2014), IL18R1-Short significantly dampened IL-18–induced NFκB activity in a dose-dependent manner across both receptor and ligand concentrations (Figure S11). Suppression exceeded 20-fold at 10 ng/mL IL-18 (adj. p = 3.14×10^-12^) and 80-fold at 100 ng/mL IL-18 (adj. p = 7.66×10^-10^) (Figure 3B). Notably, IL18R1-Short exhibited up to 30-fold more effective suppression of NFκB signaling at 100 ng/mL of IL-18 compared to the established soluble decoy IL18BP (adj. p = 1.55×10^-^⁴). (Figure 3B; Table S12).

To test whether decoy function is conserved in bat IL18R1-Short, we attempted to establish a bat-specific IL-18 reporter system. However, we were unable to purify recombinant bat IL-18, and neither human nor mouse IL-18 activated the bat receptor complex in Jamaican fruit bat primary kidney epithelial cells (Figure S12). We therefore evaluated bat IL18R1-Short in the human HEK293T reporter cell system for IL-18, where it potently suppressed NFκB activity across all IL-18 doses tested, including 1 ng/mL (adj. p = 3.69×10^-^⁸), 10 ng/mL (adj. p = 3.87×10^-10^), and 100 ng/mL (adj. p = 1.41×10^-^⁹) (Figure S8; Table S12), consistent with conserved decoy function. The mouse ortholog showed similar decoy activity in EL4 T-lymphoblasts, a mouse cell line naturally responsive to Il-18 (Figures S13). Overexpression of Il18r1-short reproducibly dampened Nfκb activity in a dose-dependent manner across both construct concentrations and Il-18 doses relative to the WT control (Figure S13).

Together, these data demonstrate that the IL18R1-Short receptor from human, mouse, and bat dampens IL-18-mediated NFκB signaling when overexpressed in mammalian cells, despite species-specific differences in the length of the L2-derived C-terminal extension—establishing membrane-anchored decoy activity as a conserved functional consequence of this ancient retrotransposon insertion.

### Mouse T cells endogenously express *Il18r1-short* that negatively regulates NFκB activity and dampens inflammatory response

We next examined whether Il18r1-short exhibits decoy activity under physiologically relevant expression levels. Mouse T cells express *Il18r1-short*, accounting for ∼20–50% of transcripts at the locus (Figure 1C; Figure 2A-D). We therefore used the mouse EL4 T-lymphoblast cells, which endogenously expresses both *Il18r1* isoforms (Table S13), to test whether Il18r1-short dampens signaling. Short-interfering RNAs (siRNAs) were designed to target the long and short isoforms independently; targeting *Il18r1-short* reduced its expression by > 70% at 24 hours compared to control cells transfected with non-targeting siRNA (Figure 3C). Consistent with a decoy receptor model, *Il18r1-short* knockdown (KD) increased Il-18–induced Nfκb activity at both 10 ng/mL (3.09-fold; adj. p = 6.69×10^-^⁴) and 50 ng/mL (2.79-fold; adj. p = 6.87×10^-^⁴), whereas *Il18r1-long* KD decreased Nfκb activity as expected (Figure 3D; Table S14). Together, these assays establish that endogenously expressed *Il18r1-short* functions as a decoy receptor of Il-18–mediated Nfκb signaling in mouse T cells (Figure 3E).

To further define the modulatory effects of Il18r1-short on downstream Il-18 induced transcriptional responses, we profiled gene-expression changes by RNA-seq of control and *Il18r1-short* KD EL4 cells at 4 hours post treatment (Figure S14-15; Table S15-17). Because a subset of genes showed baseline expression differences between conditions prior to stimulation (Figure S14), we focused subsequent analyses on genes that were comparably expressed at baseline and significantly responsive to Il-18 in both conditions, allowing a cleaner comparison of stimulus-dependent effects (Figure 4A). Most of these genes showed a larger treatment effect in *Il18r1-short* KD cells vs control (Figure 4A; Figure S15). Immune effector and inflammatory genes showed amplified Il-18–induced transcriptional responses (Figure 4B-C; Table S18-19). The most enhanced targets included *Gadd45B*, *Maf*, and *Il2rb* (p < 0.01; Figure 4C), genes linked to activated T cell Ifn-y production, effector differentiation and cytokine production, and lymphocyte proliferation and survival, respectively ^77–79^. *Clec2D*, a ligand that modulates NK and cytotoxic T cell activity ^80^, was also more strongly induced (*p* < 0.05; Figure 4C), and *Tnfsf10*, a cytotoxic ligand that promotes apoptosis ^81^, showed a similar upward trend but was not significant (Figure 4C; Table S18-19).

**Figure 4.**
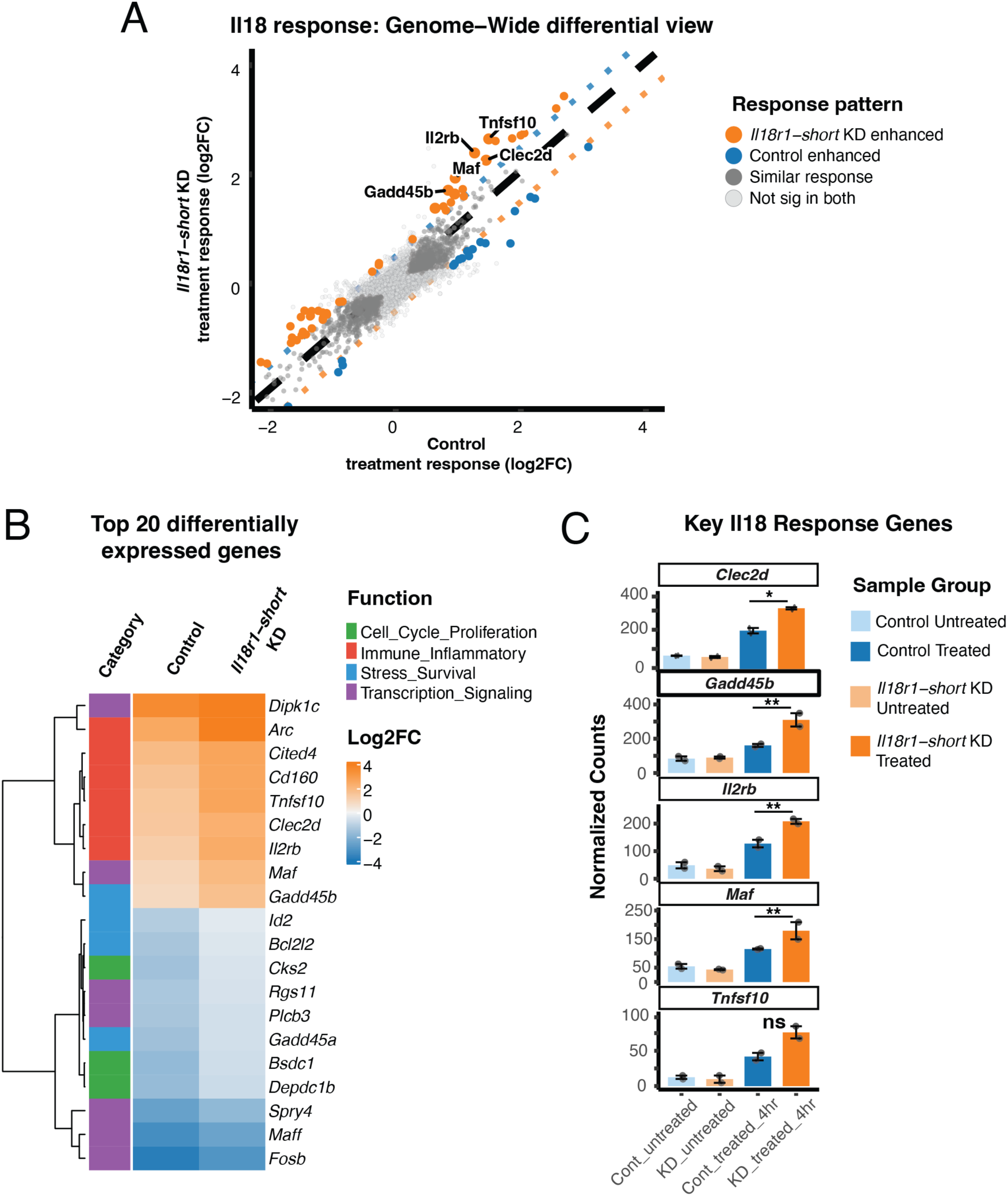
ll18r1-short dampens inflammatory responses in mouse T cells. **A** Genome-wide comparison of Il-18 treatment responses in mouse EL4 cells: control vs *ll18r1-short* KD. Orange points: genes with a stronger Il-18 treatment response in *ll18r1-short* KD cells than in control cells (Sh_FC - NT_FC > 0.5 log2FC, padj ≤ 0.05 in both conditions). Blue points: genes with a stronger 11-18 treatment response in control cells than in II18r1-short KD cells (NT_FC - Sh_FC > 0.5 log2FC, padj ≤ 0.05 in both conditions). Dark gray points indicate genes with similar responses in both conditions and light gray points indicate genes that did not reach statistical significance in either condition. Black dashed line indicates equal response; dotted lines mark ±0.5 log2FC difference thresholds. **B** Heatmap of top 20 differentially expressed genes by fold change difference. Rows represent individual genes clustered by expression pattern and assigned to a gene function category; columns show treatment-induced responses (log2FC) in control and *ll18r1-short* KD conditions. Color intensity indicates response magnitude (blue= downregulation, orange= upregulation). **C** Normalized counts for key Il18 response genes. Error bars represent± SE of the mean; n = 2 treatment replicates; individual data points overlaid. Asterisks indicate Wald test significance of differential responses: ** p < 0.01, * p < 0.05 (unadjusted p-values for hypothesis-driven genes).

A smaller subset of genes showed a larger treatment effect in control cells compared to *Il18r1-short* KD (Figure 4A, blue points), most of which were observed among genes that showed a greater downregulation in expression by 4 hours, rather than induction (Figure 4B, blue boxes). This unexpected pattern suggests that Il18r1-short silencing does not simply increase the magnitude of Il-18-responsive genes but may also reshape the broader temporal dynamics of transcriptional reprogramming during cellular stimulation. Together, these results demonstrate that Il18r1-short modulates Il-18-induced transcriptional responses in mouse T cells, with the strongest downregulation seen in immune and inflammatory gene induction.

### Transposable elements contribute to species-specific alternative transcription of immune genes

*IL18R1-Short* represents a case where species-specific co-option of an ancient retrotransposon generates an alternative immune gene transcript (Figure 1A). To assess how broadly TEs contribute to species-specific immune isoforms, we performed a transcriptome-wide intersection of predicted immune gene coding transcripts with repeat annotations for human, mouse, and Jamaican fruit bat. Considering only significantly expressed transcripts (CPM ≥ 5; ≥30% of total locus expression), we found similar numbers of immune genes expressed for each species (Figure 5A). Approximately 30% of immune genes for bat and mouse (550 for mouse; 526 for bat), and over 50% for human (905 genes), had at least one alternative isoform, indicating that alternative transcription is common at immune loci (Figure 5A). We next asked how often these alternative transcripts were predicted to remove an important protein domain. Protein domain annotation is most complete in human, so all predicted CDS across the three species were aligned to the human SwissProt database, likely increasing sensitivity for human transcripts. We found that 12% of immune genes in mouse (194), 20% in bat (277), and 40% in human (608) had at least one alternative transcript predicted to have a deleted or partially deleted protein domain (Figure 5A). Despite this quantitative difference, the most common classes of domain loss were similar across species, including disordered, intracellular, cytoplasmic, helical, signal peptide, and protein kinase domains (Figure S16). Of these domain-loss transcripts, only a small fraction was TE-derived in our analysis (Figure 5A). This is consistent with prior work showing that many TE insertions occur in UTRs where they don’t affect the CDS but can still have regulatory effects ^82–84^, not considered in this study.

**Figure 5.**
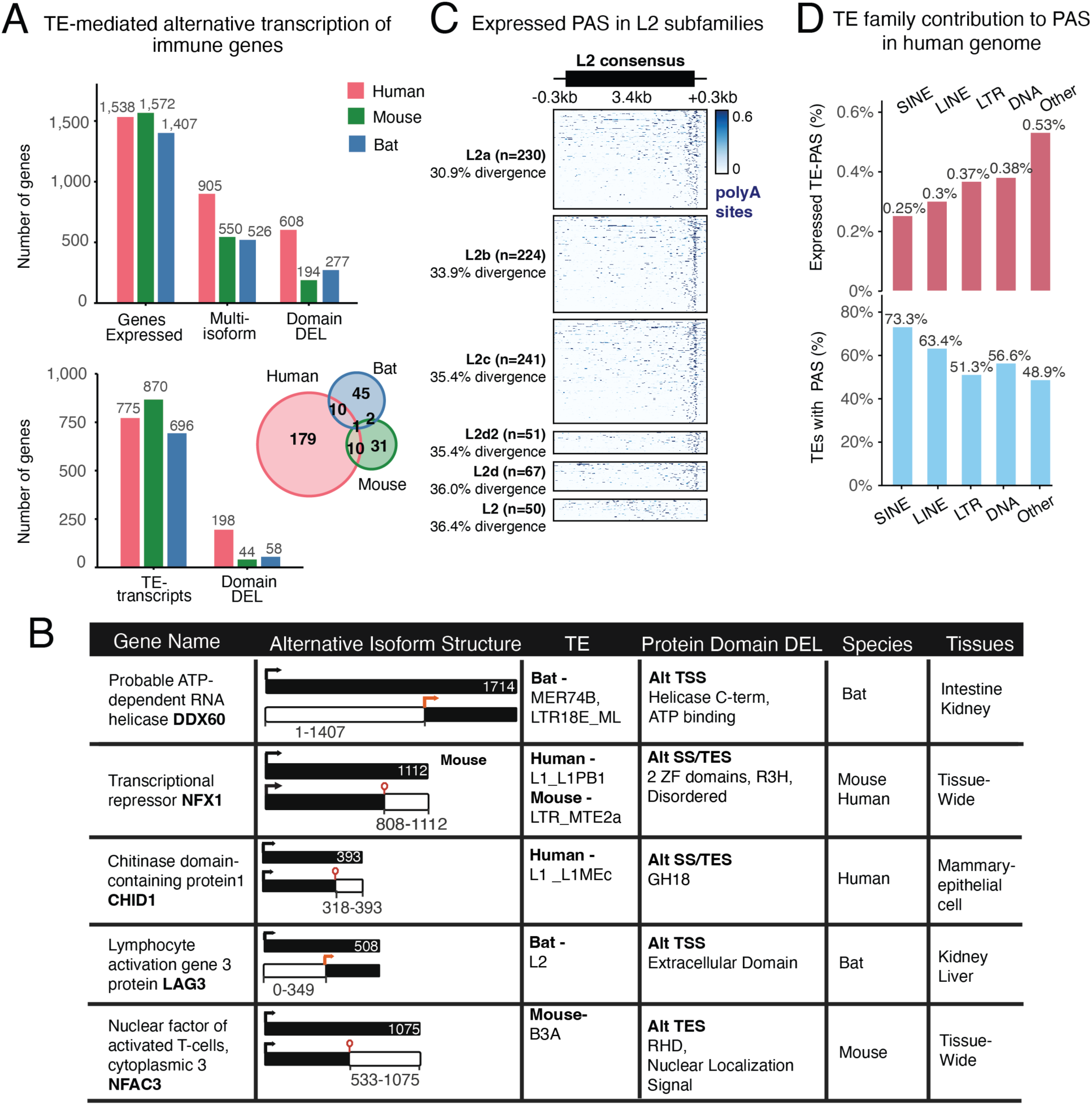
Transposable element-derived alternative transcription of immune genes across species. **A** Bar graphs showing the number of immune genes with highly expressed transcripts (CPM ≥ 5; ≥30% of locus expression) at each stage of the analysis pipeline. Top: genes with at least one expressed transcript (Genes Expressed), genes with multiple expressed transcripts (Multi-isoform), and genes with at least one transcript predicted to have a protein domain deletion (Domain DEL). Bottom: genes with at least one TE-exonization event (TE-transcripts) and the subset of those predicted to result in domain loss (Domain DEL). Venn diagram represents the subset of genes with at least one TE-exonization event resulting in domain loss. **B** Representative examples of TE-derived alternative transcripts for immune genes across species. For each gene, the full-length reference transcript (top, with total amino acid length indicated) and the alternative TE-derived isoform (bottom, with truncation coordinates corresponding to domain deletions) are shown. Red arrows indicate TE-derived alternative transcription start sites (TSS), and red circles indicate TE-derived alternative transcription end sites (TES); TE-derived splice sites noted as SS. The responsible TE family, predicted effect on protein domain structure, and species and tissues in which the alternative transcript is expressed are indicated; Tissue-wide: isoform expression in ≥ 4 tissues. **C** PAS expression signal across L2 elements in the human genome (hg38) containing expressed PAS sites (average RPM> 0.9). Each row represents one L2 element, anchored to the L2 consensus sequence (-0.3kb to +0.3kb flanking the 3.4kb element body). Subfamilies are ordered by increasing sequence divergence from their consensus. **D** Percentage of human TE insertions grouped by family harboring a PAS motif (blue) and the percentage of those with evidence of expression (Expressed PAS, red), shown for SINE, LINE, LTR, DNA, and Other repeat classes. Expression threshold: average RPM> 0.9.

Among the genes with TE-derived domain-loss transcripts, most events were species-specific (Figure 5A). *HLA-A* (MHC I) was the only gene with a domain-loss transcript identified in all three species, yet the domain deletion and the TE responsible were distinct in each case (Table S8). Beyond *IL18R1-Short*, several immune genes showed striking patterns of TE-mediated alternative transcription that are both species- and tissue-restricted, pointing to TE co-option as a source of immune gene diversification. These include *CHID1* (human mammary epithelia), *DDX60* (Jamaican fruit bat intestine and kidney), and *LAG3* (Jamaican fruit bat kidney and liver) (Figure 5B). The *LAG3* short isoform is also L2-derived, with the L2 in this case likely providing an alternative promoter rather than a PAS motif. Intriguingly, a primate-specific Alu in humans and rodent-specific B1 elements in mouse have independently inserted at the ancestral L2 promoter region of this gene, suggesting that these events abolished expression of the short isoform. For *NFX1*, the same domain truncation appears in both human and mouse but is attributed to different TE insertions (Figure 5B) in the same terminal exon region. We saw multiple TEs at the truncation site for some of these genes, like *LAG3* and *NFX1*, suggesting that certain immune loci are particularly permissive to TE insertion and have been subject to repeated, independent co-option events across lineages. Commonly, younger TEs have been implicated in species-specific adaptation, but the fact that many of our TE-derived species-specific transcripts were regulated by ancient TEs, like L2 in *IL18R1-Short*, prompted us to look closer at the age of TEs in our isoforms of interest across these three species (Figure S17-18). When we looked at the median percent divergence of all TEs in these families, we found that across species older conserved elements contribute to immune gene isoform diversity as much as younger species-specific TEs (Figure S17-18).

Finally, given that the *IL18R1-Short* PAS derives from an ancient L2 element, and that a previous study reported a truncated CD274/PD-L1 isoform similarly derived from an ancient L2 ^85^, we asked whether L2 sequences are broadly enriched for PAS motifs. Using the PolyASite database, which provides single-cell RNA-seq evidence for PAS usage, we found that all L2 subfamilies harbor a PAS motif enriched at the 3’ end of the element (Figure 5C; Figure S19). L2 elements are not, however, uniquely prone to introducing PAS motifs relative to other TE families, which is unsurprising given that PAS elements are a common feature across TE classes (Figure 5D). Notably, fewer than 1% of TE-derived PAS showed evidence of expression in the PolyASite database (average RPM > 0.9 across tissues; Figure 5D). This is consistent with the broader pattern that while TEs can seed a plethora of regulatory sequences, full co-option into stable alternative transcript expression is rare, likely dependent on both favorable genomic context at the insertion site and whether the resulting alternative transcript confers a phenotypic advantage.

## Discussion

In this study, we characterize IL18R1-Short, a transmembrane decoy receptor of IL-18 that reduces cellular responsiveness to this pro-inflammatory cytokine. We show that *IL18R1-Short* is selectively expressed across species and tissues, establishing isoform usage as a key layer of regulation for this pathway. Moreover, we provide further evidence of exonized transposable elements as a source of species-specific immune regulation ^30,85–95^.

Prior work proposed broad conservation of *IL18R1-Short* expression based on the presence of a PAS motif in intron 9 across mammals ^69^. Additionally, studies reported expression of this isoform in a limited set of mouse, rat, and human tissues using non-quantitative or low-sensitivity assays ^68–70^. We expand this framework by linking the intronic PAS to an ancient L2 insertion (∼160–186 million years old), establishing a shared cis element with the potential to regulate alternative transcription at this locus. Our isoform-resolved bulk RNA-seq analysis shows that conservation of this L2-derived PAS is necessary but not sufficient for robust *IL18R1-Short* expression. While this isoform was largely absent across mammalian species, it was highly expressed in multiple tissues in mouse and Jamaican fruit bat, with particularly strong enrichment at epithelial barrier sites, including lung, skin, and intestine (Figure 1B–C). Mouse lung single-cell data further indicate that epithelial populations account for much of the *Il18r1-short* signal. This is notable because IL-18 is more often discussed in the context of immune-cell responses, especially NK and T cell responses ^96^, whereas our data expand this view by suggesting that epithelial barriers may also be important sites of IL-18 signaling regulation. These data show that an ancient L2 insertion provides a conserved cis element whose transcriptional output varies across species, likely shaped by species- and tissue-specific locus accessibility and trans-acting regulatory factors.

Single-cell lung analysis in human and mouse also showed that the major IL-18 pathway inhibitors occupy distinct expression niches: *SIGIRR* is broadly expressed across cell types, consistent with its function as a general brake on TIR-domain signaling ^97,98^, and *IL18BP* ^99^ is also widely expressed but at much lower levels than *SIGIRR*. In contrast, *IL18R1-Short* is enriched in a narrower set of populations, including epithelial cells in mouse lung and endothelial cells in human lung. Elevated IL18R1-Short at epithelial or endothelial barriers could soak up excess IL-18 at common pathogen-contact sites and shape local effector-cell responses, analogous to decoy/scavenger systems such as D6/ACKR2 ^100,101^. A second, non-exclusive possibility is that epithelial and endothelial cells upregulate IL-18 receptor expression during inflammation and become more IL-18–responsive ^102–105^; in this context, IL18R1-Short may act as a local brake on signaling to preserve barrier integrity, similar to epithelial IL-1R2-mediated decoy regulation ^106,107^, to preserve barrier integrity. Together, these expression patterns suggest that IL-18 regulation is mediated by non-redundant inhibitors deployed in distinct cellular contexts.

Previous functional work on human IL18R1-Short had shown that over-expression of this isoform in a heterologous cell reporter system dampens NFκB activity ^70^. Using similar models, we show this extends to bat and mouse specific IL18R1-Short, despite minor differences in the 3’end of this truncated receptor across these species. In mouse T cells, *Il18r1-short* is significantly expressed, contributing to Il-18 signaling regulation. We show that silencing *Il18r1-short* expression increases NFκB activity and enhances IL18–driven immune and inflammatory gene expression, establishing the endogenous function of IL18R1-Short as a decoy that dampens pathway output.

*IL18R1-Short* is a clear example of how TE-derived regulatory sequences can introduce species-specific immune regulation. Recent lineage-specific TE insertions have often been emphasized as sources of adaptive regulatory innovation ^23,25,30,108–111^, but we show that when it comes to TE-exonization, ancient transposition events are just as likely to introduce isoforms with potential regulatory function. In general, TEs introduce a plethora of regulatory sequences, but these are mere opportunities for co-option with only a minor fraction of these events contributing to alternative host transcription. This paradigm was strikingly clear from our TE-PAS analysis, where we found that while various TE families commonly introduce PAS motifs near coding regions of the human genome, less than 1% of those sequences are expressed. One explanation is that TE insertions are subject to mutation that could disrupt their cis-regulatory potential ^108,112^. Another is that species- or tissues-specific chromatin accessibility and trans-activating factors ^113–115^ could bias the use of certain TEs over others.

Nevertheless, for disease relevant inflammatory pathways such as IL-18, the species-and tissue-restricted expression of *IL18R1-Short*—particularly in barrier tissues—underscores that baseline regulatory architecture can differ substantially across mammals. This is important to consider when extrapolating disease mechanisms and translating findings coming from research in model organisms, such as mouse, where the regulatory isoform context may be different. At the same time, these differences offer an opportunity to learn how distinct mammals tune disease-relevant inflammatory pathways, which may in turn suggest new ways to modulate these pathways therapeutically in humans. More broadly, our characterization of *IL18R1-Short* and other immune gene transcripts highlights alternative transcription as an underappreciated source of species-specific immune regulation, capable of rewiring pathway sensitivity to key signaling molecules.

Several limitations of this study warrant consideration. Our expression analysis was performed in baseline, untreated tissues; isoform ratios are likely dynamic and may shift substantially under inflammatory conditions or pathogen challenge. The single-cell data also carry caveats: the Tabula Muris and 10x mouse lung datasets showed only partial concordance in the relative abundance of *Il18r1-short* across epithelial populations, and the human 10x lung dataset had limited cell coverage, making the apparent endothelial and mast cell enrichment tentative. Finally, while we demonstrate that silencing mouse endogenous *Il18r1-short* amplifies IL-18–driven responses in T cells, the functional consequences of *IL18R1-Short* expression in primary epithelial or endothelial cells — and its impact on IL-18 signaling in the context of systemic inflammation — remain to be determined. Addressing these questions will require testing in more complex in vitro models, such as co-cultures of immune and epithelial cells, or in mouse models.

## Methods

### Primary tissue and cell collection

Two bats, one male and one female, from the Colorado State University (Fort Collins, CO) Jamaican fruit bat (*Artibeus jamaicensis)* colony were euthanized by respiratory hyperanesthesia followed immediately by thoracotomy. Tissues were aseptically removed, collected in RNAlater (ThermoFisher AM7020), and flash frozen in liquid nitrogen for subsequent RNA extraction. Additionally, we received primary Jamaican fruit bat kidney epithelial cells from the laboratory of Tony Schountz (Colorado State University) for in vitro culture experiments. These cells were originally generated from minced and trypsinized kidney tissue from *Artibeus jamaicensis* and propagated as previously described ^116^ and have subsequently been used as a cultured bat epithelial model in vitro ^117^.

Primary AT2 lung epithelial cells from mouse and human were isolated at National Jewish Health (Denver, CO) using established protocols ^118^. Mouse AT2 cells were obtained by dispase-based lung digestion and purification. Human AT2 cells were isolated from deidentified donor lungs (not used for transplantation) by perfusion and lavage, followed by elastase digestion, tissue mincing, and enrichment on a discontinuous Optiprep density gradient (1.080/1.040). Cells were further purified by IgG panning (nonadherence) and/or EpCAM (CD326) magnetic selection, as described previously ^118^. Cells were cryo-preserved or placed in RNA lysis buffer (RLT; QIAGEN 79216) for RNA extraction.

### Cell culture

Experiments were conducted in HEK293T (ATCC CRL-3216), EL4 (ATCC TIB-39), and Jamaican fruit bat epithelial kidney cells. Jamaican fruit bat epithelial kidney cells were donated by Colorado State University (Fort Collins, CO) and were derived from the same colony from which primary tissues were obtained. Cell lines were authenticated by the Molecular Biology Core Facility (Barbara Davis Center for Childhood Diabetes) at the University of Colorado Anschutz Medical Campus and periodically tested for the presence of mycoplasma using the ATCC Universal Mycoplasma Detection Kit (#30-1012K). All cell lines were maintained at 37°C and 5% CO2.

HEK293T cells were cultured in DMEM + GlutaMAX medium (ThermoFisher; 10566-024) supplemented with 10% fetal bovine serum (FBS; R&D; S11150), 2 mM L-glutamine (ThermoFisher; 25030081), and 100 U/mL penicillin-streptomycin (Gibco; 15140-122). EL4 cells were cultured in RPMI 1640 medium (Gibco; 11875085) supplemented with 10% FBS, 1× non-essential amino acids (ThermoFisher; 11140050), 1 mM sodium pyruvate (Fisher Scientific; 11-360-070), and 100 U/mL penicillin-streptomycin. Jamaican fruit bat epithelial kidney cells were cultured in DMEM + GlutaMAX medium (ThermoFisher; 10566-024) supplemented with 10% FBS (R&D; S11150), 100 U/mL penicillin-streptomycin (Gibco; 15140-122), 1× non-essential amino acids (ThermoFisher; 11140050), 1 mM sodium pyruvate (Fisher Scientific; 11-360-070), and 25 mM HEPES (ThermoFisher; 15630080). Seeding density and passaging were performed according to ATCC protocols for HEK293T and EL4 cells. Jamaican fruit bat epithelial kidney cells were maintained at 2 × 10^5^ to 5 × 10^5^ cells/mL and passaged using 0.25% trypsin-EDTA.

### Cloning of receptor constructs

A list of all plasmids used in this study is provided in Table S9; Twist oligo sequences and primer sequences for cloning are found in Table S9B. IL-18 pathway receptor CDS were obtained either by PCR amplification from cDNA or by gene synthesis (Twist Bioscience). RNA was isolated from EL4 and HEK293T cells using the Quick-RNA Miniprep kit (Zymo Research; R1054) and reverse-transcribed with LunaScript RT SuperMix (NEB; E3010L) according to the manufacturers’ instructions. Mouse IL-18 pathway receptors and human IL18BP were amplified using Q5 High-Fidelity DNA Polymerase (NEB; M0494L), whereas human IL18R1 receptors, human IL18RAP, and Jamaican fruit bat IL18R1-Short were synthesized by Twist Bioscience.

All inserts were cloned into a piggyBac overexpression vector (PB-CMV-MCS-EF1α-Puro; System Biosciences; PB510B-1) using In-Fusion cloning (Takara Bio; 638947). In-Fusion products were transformed into NEB Stable *E. coli* (NEB; C3040H) and plated on LB agar containing 100 µg/mL carbenicillin. Individual colonies were selected, plasmid DNA was purified, and construct identity was confirmed by sequencing.

For C-terminal HaloTag fusion constructs, the HaloTag sequence was amplified from pHTC HaloTag CMV-neo (Promega; G7711) using Q5 High-Fidelity DNA Polymerase and cloned into the piggyBac vector described above to generate a C-terminal HaloTag destination vector. Human IL18RAP, IL18R1-Long, and IL18R1-Short were then cloned in frame upstream of the HaloTag sequence using the same In-Fusion strategy.

### RNA-sequencing of Jamaican fruit bat tissues

Frozen tissues (intestine, kidney, heart, liver, lung, lymph node, spleen) from the Jamaican fruit bat individuals were homogenized by grinding with a pellet pestle (Fisherbrand 12-141-361) under liquid nitrogen. RNA was then extracted from ground homogenized tissues using the Quick-RNA Miniprep Kit (Zymo Research, R1054) according to the manufacturer’s protocol for tissue samples. RNA concentration was measured by Qubit, using the RNA Broad Range Assay kit (Invitrogen Q10210), and the integrity was assessed by TapeStation (Agilent; High Sensitivity RNA ScreenTape 5067-5579). All samples had RNA integrity (RIN) values between 6.6–7.2 and were used for downstream RNA-sequencing.

For short-read RNA-seq, polyA enrichment and library preparation was performed using the KAPA mRNA HyperPrep Kit (Kapa Biosystems 8098115702) according to the manufacturer’s protocols. Briefly, 500 ng of RNA was used as input, and unique dual-indexed adapters (Kapa Biosystems #KK8727) were added at a final concentration of 1.5 µM. Purified, adapter-ligated library was amplified for 11 cycles. The final libraries were pooled and sequenced on an Illumina NovaSeq X (University of Colorado Genomics Core) to obtain 20 million 150 bp paired-end reads per library. Paired-end 150bp read length FASTQ files were quality-filtered and adapter trimmed (*qin=33 ktrim=r k=31 mink=11 hdist=1 tpe tbo qtrim=r trimq=10)* using bbDuk v.38.05 ^119^; quality check was performed using FastQC v0.11.8 ^120^ and inspected through MultiQC v1.7 ^121^.

For long-read RNA-seq, cDNA libraries were prepared using the PCR-cDNA Barcoding Kit (SQK-PCB111.24; Oxford Nanopore Technologies) according to the manufacturer’s instructions. Samples were PCR-amplified with barcoded primers for 14 cycles, pooled equimolarly, and prepared for sequencing by addition of rapid sequencing adapters prior to loading. Libraries were sequenced on a R9.4.1 MinION flow cell (male bat tissues) or a R9.4.1 PromethION flow cell (female bat tissues) (Oxford Nanopore Technologies) for 72 hours. PromethION sequencing generated POD5 files while MinION sequencing generated FAST5 files, which were converted to POD5 format using the *dorado pod5 convert* function (Dorado^122^ v0.7.2). POD5 files from both bats were processed with *dorado basecaller* on super high accuracy and demultiplexed (*dorado demux with flags –not-trim, –no-classify, and –emit-fastq)*. Output FASTQ files were run through pychopper for read orientation, adapter trimming, and selection of full-length cDNA (Pychopper ^123^ v2.7.10). Pychopper was run with default parameters (pychopper *-k PCB111 -w*); output reads were combined with rescued reads and Nanofilt v2.8.0 ^124^ was used to filter reads based on a quality threshold of 10.

### Download of publicly available RNA-seq data

All RNA-seq data and accession information can be found in Table S1 and 3A. RNA-seq FASTQ files, except for ENCODE and GTEX data sets, were downloaded from public repositories using fasterq-dump from the NCBI SRA toolkit v3.0.2. GTEx V9 Protected Access long-read RNA-seq FASTQ files (https://gtexportal.org/home/protectedDataAccess) were downloaded on April 2023 using AnViL (https://anvil.terra.bio/). Long-read RNA-seq BAM files were downloaded from the ENCODE portal (https://www.encodeproject.org) on March 2023 for *Homo sapiens* long-read RNA-seq experiments (i.e. using the following search: https://www.encodeproject.org/search/?type=Experiment&control_type!=*&assay_term_name=long%20read%20RNA-seq&replicates.library.biosample.donor.organism.scientific_name=Homo%20sapiens&status=released). BAM files were downloaded using wget, then sorted and indexed using samtools v1.8 ^125^. Files were renamed to include tissue or cell line identifiers.

### Jamaican fruit bat transcriptome assembly from RNA-seq

Processed long-reads were aligned to the CSHL GCF_021234435.1 genome assembly using minimap2 ^126^ (*-x splice*). Output SAM files were converted to BAM, filtered for alignment quality of 10 or higher, and merged, sorted, and indexed with samtools v1.8. This merged BAM file was used as input for long-read based transcriptome assembly carried out by Stringtie v2.2.1 ^127^ (*-L --conservative*). The output Stringtie GTF file was merged with the reference RefSeq GTF file for GCF_021234435.1, and redundant and single-exon transcripts were removed (gffcompare *-M;* GffCompare v0.12.9 ^128^).

### Isoform-aware quantification of *IL18R1*

Species-specific RefSeq GTF files were obtained from UCSC for all mammalian genomes (Table S6) except human, mouse, and Jamaican fruit bat. For the Jamaican fruit bat, our long-read generated GTF was used (GSE324647); for human, the GTEx V9 FLAIR transcriptome was used (https://www.gtexportal.org/home/datasets); for mouse, the TAMI long-read transcriptome (https://vollmerslab.soe.ucsc.edu/tami/) was merged with the GENCODE mouse annotation (https://ftp.ebi.ac.uk/pub/databases/gencode/Gencode_mouse/release_M37/gencode.vM37.annotation.gtf.gz) using GffCompare, and redundant and single-exon transcripts were removed.

For short-read quantification, each GTF file was modified to retain one long and one short *IL18R1* isoform. As most species lacked annotation for the short isoform, it was generated using exons 1–9 of *IL18R1-Long*, with exon 9 extended to the end of the L2-derived PAS sequence; modified GTFs can be found in the Zenodo project for this study (10.5281/zenodo.19022336) and the *IL18R1* isoform IDs are provided for each species (Table S6). GTF files were converted to FASTA format using gffread (Cufflinks ^129^ v2.2.1) to generate a salmon transcript index. Downloaded FASTQ read files were quality filtered and adapter trimmed as previously described for Jamaican fruit bat generated short-read RNA-seq. Quantification was performed using salmon ^130^ v1.10.0 with parameters *--libType A -p 8 --rangeFactorizationBins 4 --numBootstraps 10000*. TPM values for *IL18R1* isoforms from mammalian short-read RNA-seq are found in Table S1.

For long-read quantification, bambu ^131^ v3.4.1 (*se.quantonly, discovery = FALSE)* was used. For human, mouse, and Jamaican fruit bat, all isoforms detected at the *IL18R1* locus were included to enable unbiased quantification (Table S3B). A description of all isoforms and their coding sequences is provided in Figure S4. Only isoforms with detectable expression (CPM ≥ 1) were considered; CPM values from isoforms sharing the same intact coding sequence were summed to obtain total expression for *IL18R1-Long* and *IL18R1-Short* (Table S3A).

### Single Cell RNA-seq

Mouse lung scRNA-seq data were downloaded from ArrayExpress (E-MTAB-13275, ERR11871099–104), human lung data from dbGaP (phs001750, SRR8295001, SRR8295012, SRR8295023, SRR8295034, SRR8295053, SRR8295056–57, SRR8295060), and Jamaican fruit bat intestine data from GEO (GSE243982, SRR26172138–43, SRR26172154–59, SRR26172164–69).

Illumina adapter and polyA/polyG sequences were trimmed with bbDuk (bbMap *ktrim=l k=23 mink=11 hdist=1 skipr1*). Trimmed reads were aligned with STARsolo ^132^ (v2.7.11B) to hg38 (human), mm39 (mouse), or GCF_021234435.1 (Jamaican fruit bat), using chemistry v2 or v3 10X Genomics cell barcodes appropriate to the dataset and standard solo parameters for CB/UMI demultiplexing, EmptyDrops_CR cell filtering, and EM multi-mapper handling. Resulting BAM files were filtered to remove multimapping reads with samtools (v1.16.1). Cell filtering, normalization, and clustering were performed in Scanpy ^133^ (v1.11.4, Python v3.11.3, AnnData ^134^ v0.12.2). Cells with fewer than 100 detected genes and genes present in fewer than 3 cells were excluded. Counts were normalized to 10,000 per cell, and the top 2,000 variable genes were used for principal component analysis (PCA) and UMAP visualization. Leiden clustering ^135^ was performed across multiple resolutions and visually inspected for cell type annotation. Human and mouse cell types were annotated using CellAnnotator ^136^ (v0.2.1) with the OpenAI model gpt-4o-mini; bat clusters were annotated manually using marker genes, with ambiguous clusters resolved by querying GPT-4 with the top 7–10 differentially expressed genes. Cells labeled as “Unknown” or remaining ambiguous were excluded from isoform analyses and visualization.

*IL18R1* isoform expression was quantified by defining 501 bp windows centered on each isoform’s polyadenylation site (300 bp upstream to 200 bp downstream). Reads overlapping these windows were counted with featureCounts ^137^ (Subread v1.6.2) and UMI-tools ^138^ count (v1.1.6; *--per-gene --per-cell;*), then normalized to 10,000 reads per cell alongside detected genes. Isoform-specific counts were averaged per cell type and cell detection rates were reported. Differential isoform usage plots were generated in R (v4.3) with ggplot2 (v4.0.1).

### HaloTag (pHTC) Imaging of Human IL-18receptors

HEK293T cells were transfected in separate reactions with 3 µg of human IL18R1-Long-pHTC, IL18R1-Short-pHTC, or IL18RAP-pHTC using FuGENE 4K (Promega; E5911) at a 3:1 reagent:DNA ratio. After 24 h, 10,000 cells were seeded into fibronectin-coated 96-well SensoPlates (Greiner Bio-One; 655892; 1 mg/cm^2^). After an additional 24 h, cells were incubated with 200 nM JFX-549 ligand (Promega; HT1020) for 1 h at 37°C. For IL-18 treatment conditions, cells were exposed during the final 30 min of this incubation to either 0 or 100 ng/mL human IL-18 (InvivoGen; rcyec-hil18) in medium containing 200 nM JFX-549.

Cells were then washed with 1× PBS and fixed in 4% paraformaldehyde for 15 min at room temperature, followed by three 5-min washes in 1× PBS. Cell membranes were stained with wheat germ agglutinin-Alexa Fluor 488 and nuclei were stained with DAPI, each for 10 min with PBS washes in between. Cells were washed again and imaged immediately.

Cells were imaged using the NIS Elements imaging software (v5.42.03) on a Nikon Spinning Disc Confocal Yokogawa CSU X1. Images were taken in channels 405 nm, 488 nm and 561 nm at an exposure of 300 ms. The laser intensity was 20% for the 561 nm channel and 10% for the 405 and 488 nm channels. Images were analyzed using Fiji-ImageJ (v 2.9.0 / 1.53t) ^139^.

### Luciferase reporter assay measuring NFκB activity

NFκB activity was measured using the Nano-Glo Dual-Luciferase Reporter Assay System (Promega; N1620) in HEK293T, EL4, and Jamaican fruit bat epithelial kidney cells.

For HEK293T assays, cells were transfected with FuGENE 4K (Promega) at a 3:1 reagent:DNA ratio using 802 ng total plasmid DNA per condition, consisting of 2 ng Firefly control vector (pGL4.50[luc2/CMV/Hygro], Promega), 100 ng human IL18R1-Long, 100 ng human IL18RAP, 100 ng NFκB NanoLuc reporter vector (Promega; N1111), and 500 ng of human IL18R1-Short, Jamaican fruit bat IL18R1-Short, or human IL18BP. After 24 h, cells were seeded at 10,000 cells per well in opaque-bottom 96-well plates (Greiner Bio-One) in medium supplemented with 2 µg/mL puromycin and 50 µg/mL hygromycin. After an additional 24 h, cells were treated with 0, 1, 10, or 100 ng/mL human IL-18 (InvivoGen; rcyec-hil18) for 4 h at 37°C before luciferase measurement.

For EL4 knockdown assays, cells were transfected by NEON electroporation (Invitrogen) in 10 µL reactions (1200 V, 20 ms, 2 pulses) with 100 ng Firefly control vector and 500 ng NFκB NanoLuc reporter vector (Promega; N1111). For siRNA knockdown experiments, custom siRNA oligonucleotides were obtained from Horizon Discovery (Table S10), and 500 nM Il18r1-short-targeting, Il18r1-long-targeting, or non-targeting control siRNA was introduced during electroporation. At 48 h post-transfection, cells were treated with 0, 10, or 50 ng/mL recombinant mouse Il-18 (R&D Systems; 9139-IL-050) for 4 h. Following treatment, 50,000 cells from each condition were seeded into opaque-bottom 96-well plates for luciferase measurement.

For EL4 overexpression assays, cells were transfected by NEON electroporation under the same conditions with 100 ng Firefly control vector, 500 ng NFκB NanoLuc reporter vector, and either 500 ng or 1 µg Il18r1-short overexpression plasmid. An mCherry overexpression plasmid was added as needed to equalize total DNA input across conditions. At 48 h post-transfection, cells were treated with 0, 10, or 100 ng/mL recombinant mouse Il-18 for 4 h, and 50,000 cells from each condition were then seeded into opaque-bottom 96-well plates for luciferase measurement.

For Jamaican fruit bat epithelial kidney cell assays, cells were transfected with FuGENE 4K (Promega) at a 3:1 reagent:DNA ratio using 2 µg total plasmid DNA per reaction, consisting of 100 ng Firefly control vector, 400 ng NFκB NanoLuc reporter vector, and 300 ng each receptor plasmid. Receptor combinations included bat IL18R1-Long plus bat IL18Rap, human IL18R1-Long plus human IL18RAP, or the mixed-species combinations shown in Figure S12. An mCherry control plasmid was added as needed to equalize total DNA input. After 24 h, 30,000 cells were seeded into opaque-bottom 96-well plates (Greiner Bio-One). After an additional 24 h, cells were treated with 0 or 100 ng/mL human IL-18 (InvivoGen; rcyec-hil18) or mouse Il-18 (R&D Systems; 9139-IL-050) for 4 h at 37°C before luciferase measurement.

For luminescence detection, ONE-Glo reagent was added to each well, plates were shaken at 300–600 rpm for 10 min, and Firefly luminescence was measured on a SpectraMax iD3 plate reader (Molecular Devices). NanoDLR Stop & Glo reagent was then added, plates were shaken for 10 min, and NanoLuc luminescence was measured. NFκB reporter activity was calculated as NanoLuc luminescence normalized to Firefly luminescence. For EL4 experiments, normalized values were additionally expressed as fold change relative to untreated controls. Statistical significance was assessed using unpaired two-sided t-tests.

### Quantitative RT-PCR

Gene expression was quantified by one-step RT-qPCR using the Luna Universal One-Step RT-qPCR Kit (New England Biolabs; E3005L) according to the manufacturer’s instructions. Each 10-µL reaction contained 25 ng RNA and 0.4 µM of each primer (Table S9) and was run on a CFX384 Touch Real-Time PCR Detection System (Bio-Rad) using the following cycling conditions: 55°C for 10 min, 95°C for 1 min, followed by 39 cycles of 95°C for 10 s and 60°C for 30 s, with a final step at 60°C for 30 s. Amplification specificity was confirmed by melt curve analysis. Technical duplicates were averaged before downstream analysis. Target gene Cq values were normalized to the housekeeping gene *CTCF*.

For mouse and human AT2 cells, relative expression was calculated using the 2^(-ΔCq) method, and data are presented as mean ± s.e.m. of biological replicates (Table S11). For EL4 siRNA knockdown experiments, fold change was calculated using the 2^(-ΔΔCq) method relative to the non-targeting siRNA control, and data are presented as mean ± s.e.m. of biological replicates (Table S13).

### RNA-seq of Il-18 treatment course in EL4 cells following *Il18r1-short* knockdown

EL4 cells were transfected by NEON electroporation, as described above, with 500 nM Il18r1-short-targeting siRNA or a non-targeting siRNA control. At 48 h post-transfection, cells were treated in duplicate with 0 or 10 ng/mL recombinant mouse Il-18 for 4 h. Approximately 1 × 10^6 cells per condition were then collected in DNA/RNA Shield (Zymo Research; R1100-50-E) and submitted for short-read RNA-seq (Plasmidsaurus; Illumina NovaSeqx single-end; oligo-dT priming), yielding 10–15 million reads per sample.

### Differential gene expression of EL4 Il-18 treatment course

RNA-seq data was downloaded from Plasmidsaurus as deduplicated BAM files. Gene-level counts were quantified using featureCounts (v2.0.1) with the parameters *-T 1 -O -p -s 2 -t exon -g gene_name*. Differential expression analysis was performed in DESeq2 ^140^ (v1.42.0). The experimental design included two knockdown conditions (non-targeting control and *Il18r1-short* knockdown) and two treatment states (untreated or Il-18 treated for 4 h; n = 2 biological replicates per group). An interaction model was fit (*design = ∼ condition + KD + condition:KD*) to assess whether knockdown modified Il-18 responsiveness. Treatment-induced fold changes for each condition were extracted independently. Log2 fold changes were subjected to shrinkage estimation using apeglm ^141^ (v1.24.0) for model coefficients and ashr ^142^ (v2.2-54) for custom contrasts.

Genes passing quality filters (significant in both conditions with p_adjusted < 0.05, no baseline expression differences with |log2FC| < 1.0 and p_adjusted > 0.1, and ≥20 normalized counts in treated samples) were ranked by fold change difference magnitude for visualization (Table S18). For validation of key Il-18 pathway genes (Il2rb, Tnfsf10, Clec2D, Gadd45B, and Maf), statistical significance of fold change differences was assessed using Wald tests, where *Z = (Log2Fold_Il18r1sh_KD - Log2Fold_NT) / √(SE_Sh^2^ + SE_NT^2^)*. Unadjusted p-values are reported for these hypothesis-driven genes (Table S19). Functional grouping was performed using Gene Ontology annotations. Figures were generated using ggplot2 (v3.4.2) and ComplexHeatmap ^143^ (v2.18.0).

### TE-derived alternative transcripts of immune genes with protein domain deletions

Long-read GTFs for the human, mouse, and Jamaican fruit bat were converted into FASTA format using gffread (Cufflinks v2.2.1). ORFs were identified with TransDecoder LongOrfs (v5.5.0) and were annotated against the human SwissProt ^144^ databse (October 2024) using BLASTp ^145^ (v.2.7.1). TransDecoder was then run with the *Predict* option and BLASTp outfmt-6 file to identify the single best coding ORF for each transcript. Species-specific peptide sequences were matched to their corresponding human peptide sequences, using the BLASTp output, and aligned with MUSCLE ^146^ (v3.8). Alignment gaps >10 aa were classified as deletions, and gaps within 15 aa of each other were merged to facilitate downstream domain assignment.

The resulting BED file contained alignment gap coordinates, and the corresponding species-specific transcript ID. Uniprot domain files were downloaded from UCSC (https://hgdownload.soe.ucsc.edu/gbdb/hg38/uniprot; October 2024) as bigBED files and converted to BED (UCSC toolkit ^147^) format for intersection with alignment gaps BED file generated in the previous step. Domain annotations included both SwissProt-reviewed and TrEMBL-computational entries; only SwissProt-reviewed annotations were retained for analysis. Bedtools ^148^ (v.2.28.0) was used to intersect domain files (Cytoplasmic, Extracellular, Signal (Peptide), Transmembrane, (others of) Interest) with alignment gaps to identify truncated or deleted domains. Intersections were then filtered to retain larger, likely disruptive deletions, using the following minimum sizes: >50 aa for Cytoplasmic, Extracellular, and other features of interest; >20 aa for Signal peptide; and >15 aa for Transmembrane domains. Filtered outputs were merged into a final table of domain deletions. This table was further restricted to significantly expressed transcripts (CPM >5 and isoform usage >30%) from immune genes (GO: 0006955, 0045088, and 0006954) to identify candidate events of interest.

For TE-exonization analysis, we used repeat annotation from Dfam.org for human (hg38; hg38.dfam.ucsc.filtered.bed) and mouse (mm39; mm39.dfam.ucsc.filtered.bed), downloaded from UCSC. For the Jamaican fruit bat we merged UCSC repeat database (GCF_021234435.1; https://ftp.ncbi.nlm.nih.gov/genomes/all/GCF/021/234/435/GCF_021234435.1_CSHL_Jam_final/GCF_021234435.1_CSHL_Jam_final_rm.out.gz) with a custom bat-specific library ^149^ using bedtools merge (*-i - -s -c 4,6 -o distinct,distinct*; v.2.28.0). Exons from significantly expressed transcripts were classified as internal or external, and exon boundary coordinates were extracted. Internal exon boundaries were intersected with repeat annotations using default 1-bp overlap criterion, whereas external exon boundaries were intersected using bedtools intersect *-f 0.1* within a 100-bp window of the 5′ or 3′ transcript boundary. To identify transcripts in which TE intersections mapped to deleted domains, TransDecoder GFF3 output was used to project alignment gap peptide coordinates back to genomic coordinates and compare them with TE intersection coordinates. Final TE-derived alternative transcripts are listed in Table S20.

### Identification of TE-derived polyadenylation sites

Repeat annotation for the human genome (hg38; hg38.dfam.ucsc.filtered.bed) was filtered to remove TEs on unplaced or random contigs. PAS annotations were downloaded from the PolyASite 3.0 atlas (atlas.clusters.3.0.GRCh38.GENCODE_42.bed from https://polyasite.unibas.ch/atlas_sc), which provides per-site average expression scores derived from single-cell 3’ sequencing data. The atlas BED file was converted to a bigWig by extracting the average RPM column, merging overlapping sites by taking the maximum score, and converting to bigWig format using bedGraphToBigWig from the UCSC toolkit. TE loci were overlapped with PAS sites using Bedtools (v2.28.0) with a 10bp window, and PAS-containing TEs were classified into five supergroups (LINE, SINE, LTR, DNA, Other) based on their Dfam repeat annotation. Sites were filtered for expression using a minimum average RPM score of 0.9, yielding 9,673 expressed TE-derived PAS loci from a total of 5,017,430 annotated TE loci.

### L2 subfamily analysis and visualization of polyadenylation sites

L2 elements (including subfamilies L2, L2A, L2B, L2C, L2D, L2D2) overlapping expressed PAS sites were identified for each subfamily using Bedtools (v2.28.0), allowing a 10bp window. For visualization, each L2 fragment was expanded to its theoretical full-length coordinates using positional information from the Dfam RepeatMasker output file (hg38.dfam.fa.out), which records the position of each genomic fragment relative to the ∼3.4kb L2 consensus sequence. Expanded L2 elements were visualized using Deeptools ^150^ (v3.0.1). The Deeptools computeMatrix function (reference-point mode, anchored at TSS, *window - 300bp to +3,700bp, binSize=1Bp*) was first used to calculate the matrix. The Deeptools plotHeatmap function was then used to plot signal from the PolyASite RPM bigwig for each L2 element. In a separate analysis, FIMO v5.1.0 (part of the MEME Suite ^151^ for Motif Analysis) was used to scan the hg38 genome for canonical motifs (AATAAA) at a p-value threshold of 1×10^-3^, and the resulting motif bigWig was visualized across all L2 elements using the same deeptools workflow (Figure S19).

## Supplemental Information

Document S1 contains Figures S1-S19. Supporting evidence for Results section. SuppTables file contains Tables S1-S20. Supporting evidence for Results section.

## Acknowledgements

We thank the BioFrontiers Institute at the University of Colorado Boulder for computational and imaging support, including the BioFrontiers IT department and the Advanced Light Microscopy Core (RRID: SCR_018302). Imaging was performed on a spinning disc confocal Nikon Ti-E microscope supported by the BioFrontiers Institute and the Howard Hughes Medical Institute. We thank Dr. Coby McDonald (formerly at Colorado State University, now at the U.S. Fish and Wildlife Service) for the Jamaican fruit bat tissues extractions and the preparation of the short-read RNA-seq libraries, the University of Colorado Anschutz Genomics Core for sequencing support, and the Donor Alliance of Colorado and IIAM for assistance in procuring human lung tissues used for AT2 isolation in this study.

## Declaration of AI-assisted technologies

The authors declare that OpenAI model gpt-4o-mini was used for single cell annotation, after which the authors reviewed and edited the content as needed. Claude Opus 4.5 was used to help generate code for identifying TE-derived transcripts and for statistical analysis of differential gene expression in EL4 cells, with author guidance and verification at every step.

## Data Availability

Raw and processed sequencing data generated in this study have been deposited in NCBI under BioProject accessions **PRJNA1434542** and **PRJNA1434540**. Gene expression matrices, processed transcript quantifications, and related expression files have been deposited in GEO under accessions **GSE324648**, **GSE324647**, and **GSE324646**. Custom scripts used for data processing and analysis are available on GitHub at: https://github.com/ordoneza2025/Long-Read-RNAseq-Analysis---ONT https://github.com/ordoneza2025/IL18R1-isoform-quantification https://github.com/ordoneza2025/Alternative-transcripts-domain-deletions https://github.com/atmaivancevic/polyAsite-analysis

Annotation files, supporting reference files, and additional processed outputs used for alternative isoform analysis and quantification in this study are available on Zenodo at https://doi.org/10.5281/zenodo.19022336

