## Supplementary figures S1-S19 for "An ancient retrotransposon provides species-specific tuning of IL-18 inflammatory signaling"

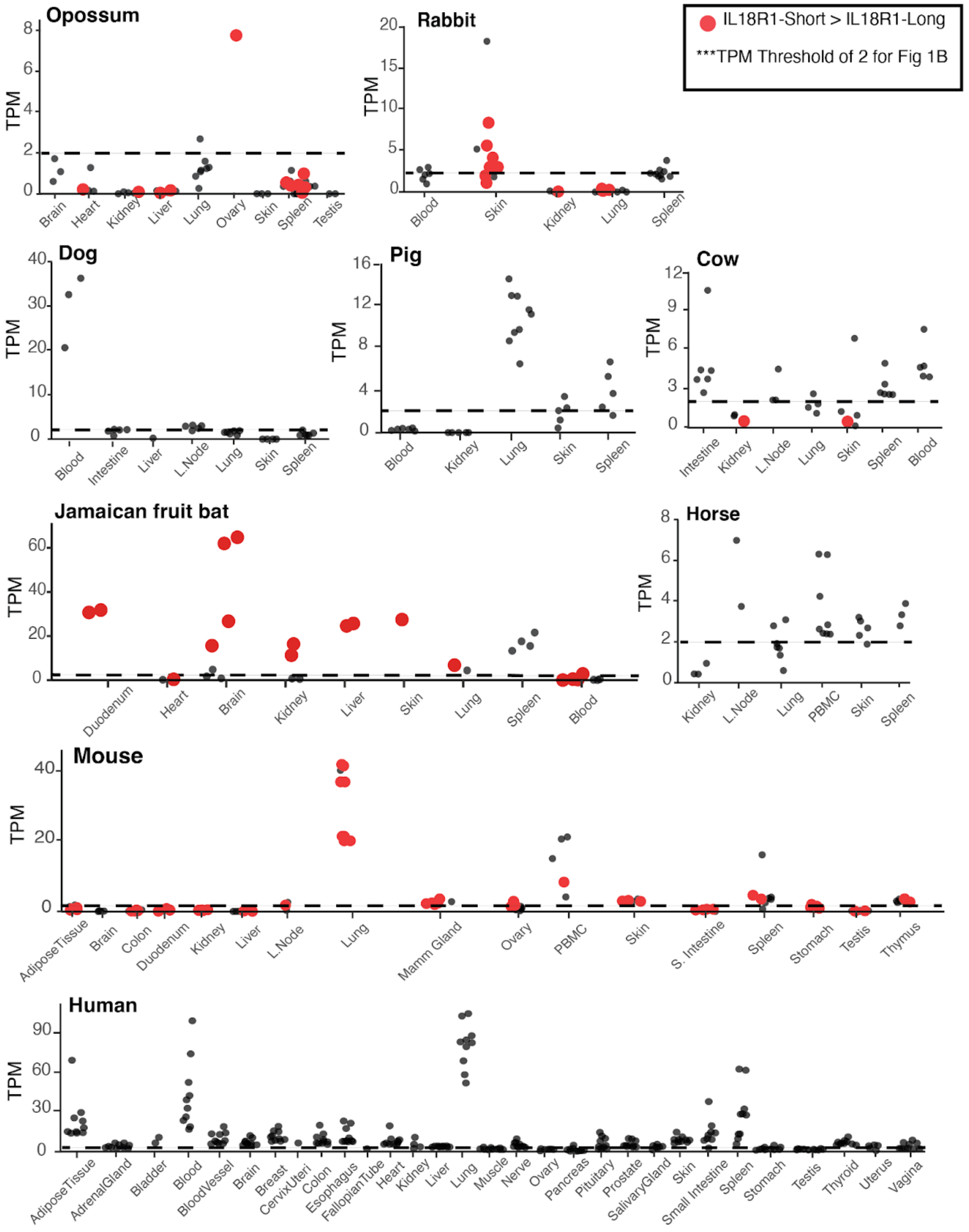


**Fig. S1. *IL18R1* expression across mammals and tissues.** *IL18R1* locus expression quantified from short-read RNA-seq data is plotted for every species and tissues assayed; data is separated by species. A threshold of 2 TPM was set to indicate significant expression for the locus (dashed line), and this data was used for **Figure 1B.** Red data points represent samples where *IL18R1-Short* expression was higher than *IL18R1-long*. PBMC = Peripheral Blood Mononuclear Cells; L. Node = lymph node; S. Intestine = small intestine.

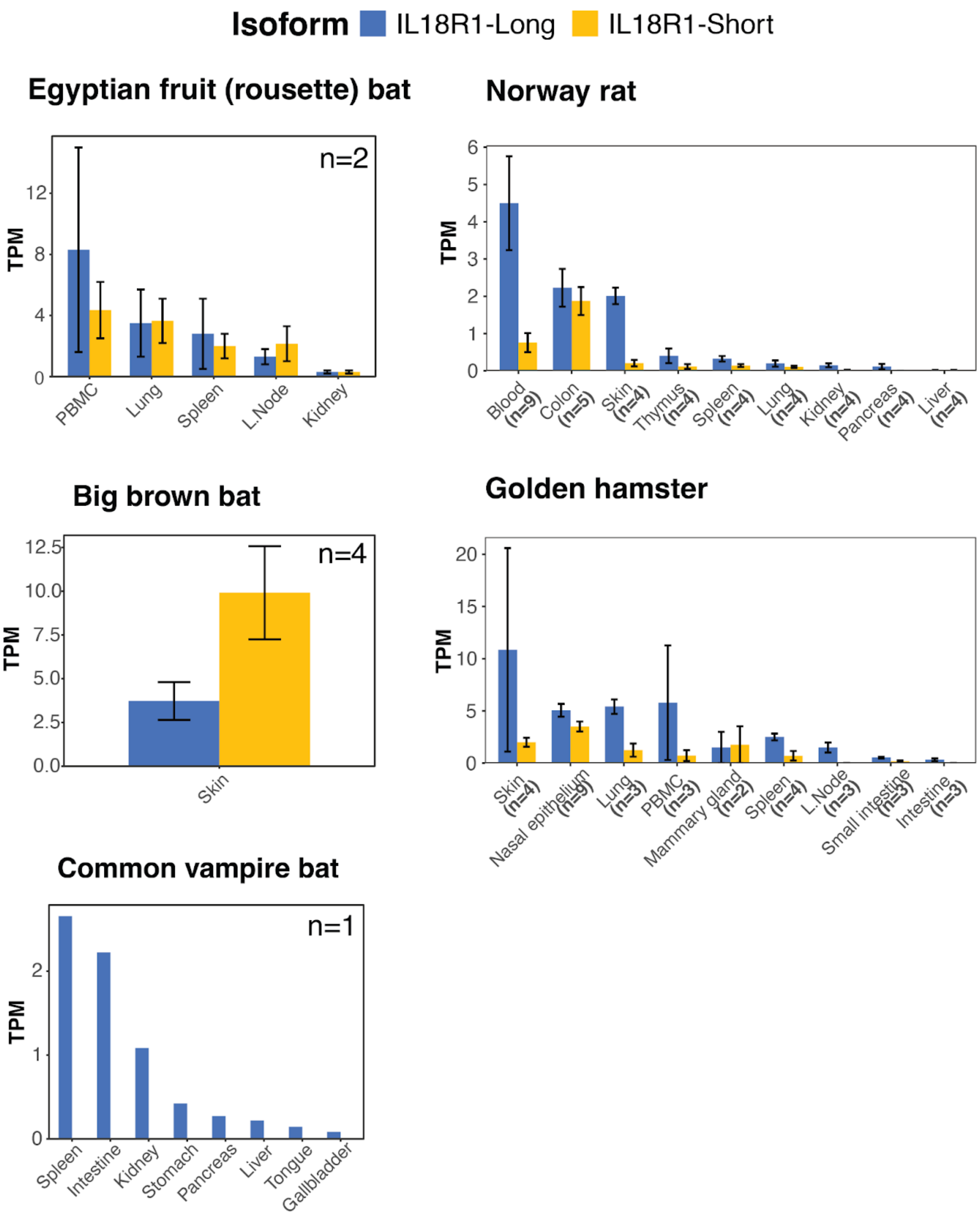


**Fig. S2. *IL18R1* expression in other bats and rodents.** Bar graphs showing average *IL18R1* isoform expression in primary tissues of other bats (left side graphs) and rodents (right side). *IL18R1-Long* is shown in blue and *IL18R1-Short* is shown in yellow. Error bars represent ± SE of the mean. This is supplementary information for **Figure 1 B-C.**

### IL18R1 isoforms detected from long-read RNAseq

| ID | CDS | Notes | Expression |
|---|---|---|---|
| ENST00000233957.5 | IL18R1_ Long | Intact CDS | ✓ |
| 2d3d1bbc-01c2-4183-aa57-854d90e4e9ca_ ENSG00000115604.10 | IL18R1_ Short | Intact CDS; Del in 5'UTR | ✓ |
| 5e1f21b1-1af6-4760-b85b-b4379ec62d74_ ENSG00000115604.10 | IL18R1_ Short | Intact CDS | ✓ |
| 9407a40a-6d52-4e9d-8849-bc526ceb5fc7_ ENSG00000115604.10 | IL18R1_ Short | Early frameshift due to exon skipping. | × |

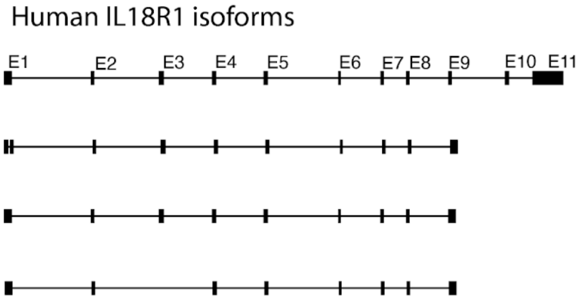


| ID | CDS | Notes | Expression |
|---|---|---|---|
| TCONS_ 00001115 | IL18R1_ Long | Intact CDS | ✓ |
| TCONS_ 00001112 | IL18R1_ Short | Intact CDS | ✓ |
| TCONS_ 00001113 | IL18R1_ Short | Intact CDS; Del in 5'UTR | ✓ |
| TCONS_ 00001114 | IL18R1_ Short | Early stop codon | × |

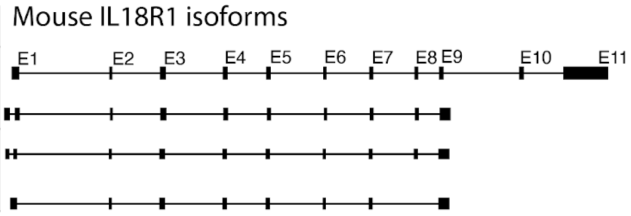


| ID | CDS | Notes | Expression |
|---|---|---|---|
| TCONS_ 00026379 | IL18R1_ Long | Intact CDS | ✓ |
| TCONS_ 00026380 | IL18R1_ Long | Del_316_A | × |
| TCONS_ 00025744 | IL18R1_ Long | Ins_474-476_GKK | × |
| TCONS_ 00025745 | IL18R1_ Long | Del_422-474 TIR Domain | AJ1_Lung |
| TCONS_ 00026381 | IL18R1_ Long | Ins_474-476_GKK Del_316_A | × |
| TCONS_ 00025743 | IL18R1_ Long | Intact CDS Del in 5'UTR Truncated 3'UTR | ✓ |
| TCONS_ 00025740 | IL18R1_ Short | Intact CDS | ✓ |
| TCONS_ 00025739 | IL18R1_ Short | Intact CDS; Del in 5'UTR | ✓ |
| TCONS_ 00025742 | IL18R1_ Short | Del in 5'UTR Del_316_A | × |
| TCONS_ 00025741 | IL18R1_ Short | Del_316_A | × |

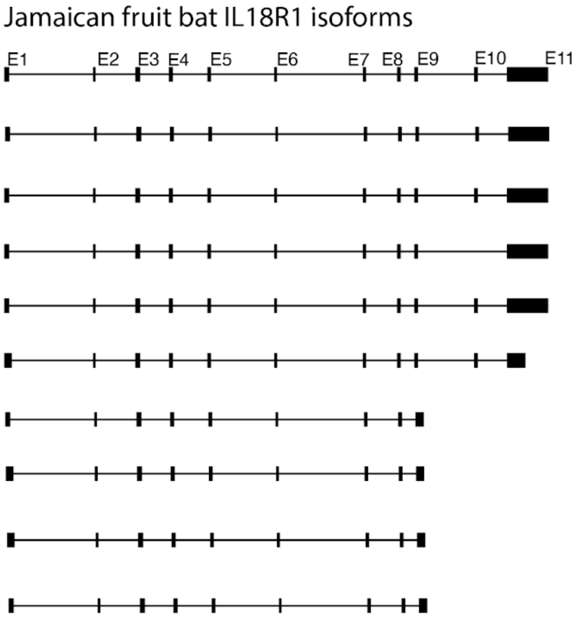


**Fig. S3. Annotated *IL18R1* isoforms from long-read cDNA.** Schematic representation of all *IL18R1* isoforms detected from long-read RNA-seq in human, mouse, and Jamaican fruit bat. Tables provide transcript ID from GTF files (Zenodo), description of the predicted CDS, and expression level for each isoform (details in table S3B). *IL18R1-Long* and *IL18R1-Short* isoforms with intact CDS and CPM ≥ 1 were considered for quantification in **Figure 1C**.

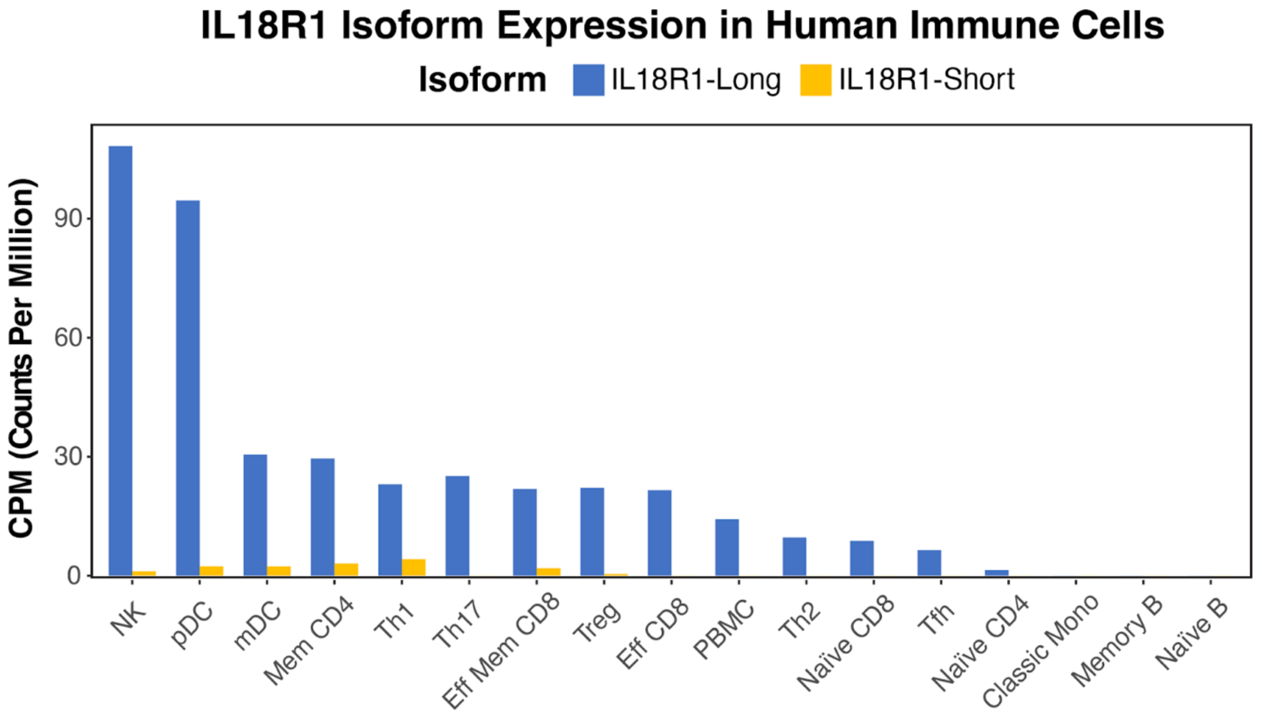


**Fig. S4. *IL18R1* expression in human immune cells.** *IL18R1* expression (CPM) from long-read RNA-seq of isolated human immune cell populations. Quantifications for these samples and full names for cell types are found in Supplementary Table 3A. A subset of this data is included in **Figure 1C.**

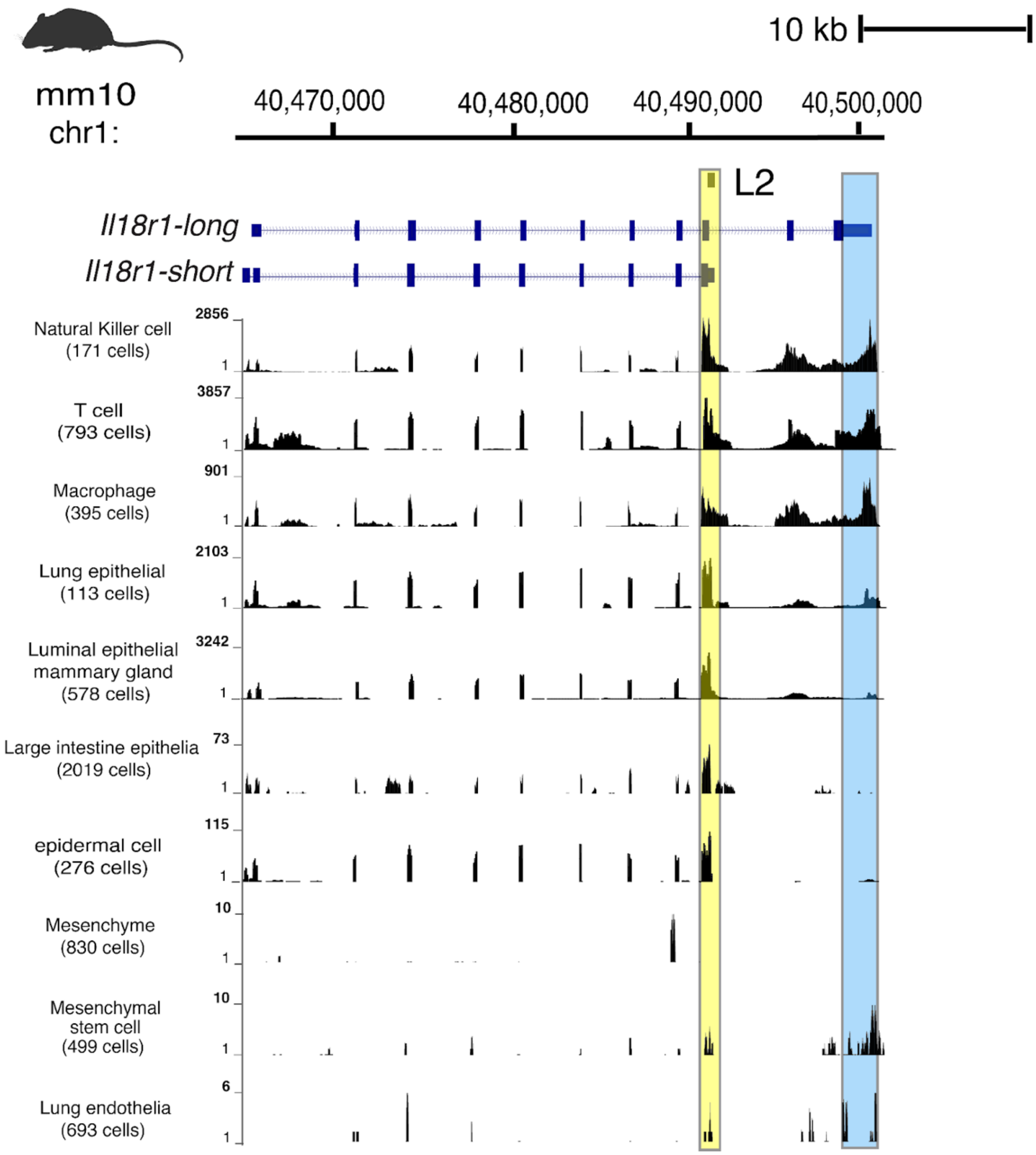


**Fig. S5. Single-cell RNA-seq of mouse tissues at the *Il18r1* locus.** Bigwig alignments (mm10 mouse genome assembly) of *Il18r1* locus expression from UCSC Tabula Muris single cell data track; 20 organs - Smart-seq2 and droplet data. Isoform maps for *Il18r1* are shown. Read depth scale (1 to indicated maximum) is shown for each population; cell numbers per population are indicated in parentheses; canonical transcriptional end site is highlighted in blue and L2-derived truncation is highlighted in yellow. A subset of this data is included in **Figure 2A.**

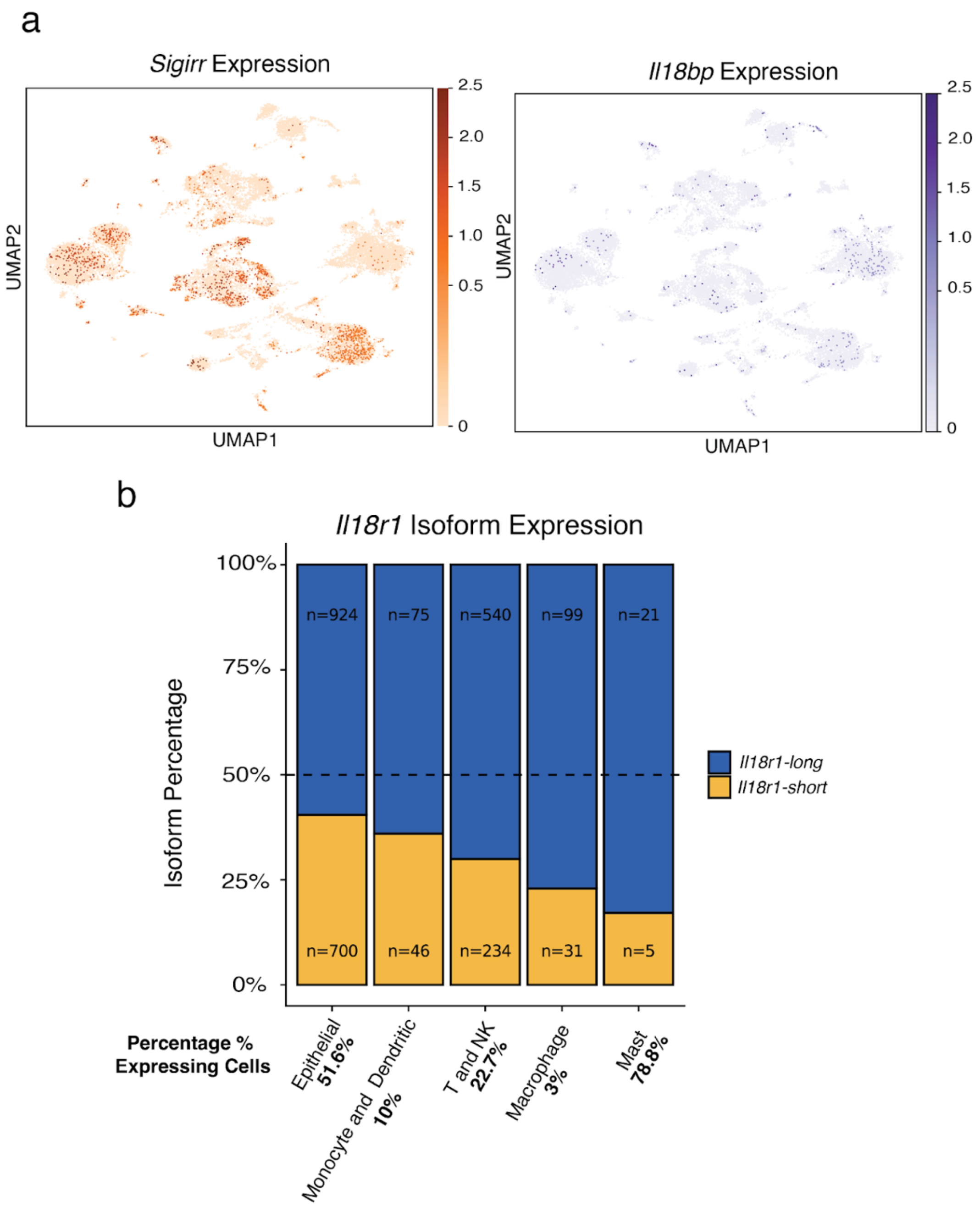


**Fig. S6. Single-cell RNA-seq of mouse Il18 regulators in the lung. A** UMAP plots showing normalized expression of mouse *Sigirr* and *Il18bp* across all profiled lung cells; scale bar represents (log1p (counts per 10K). Cell population UMAP is found in **Figure 2B** and counts for these genes are included in the dot blot in **Figure 2D**. **B** Stacked bar plots showing the average percent expression for *Il18r1* isoforms from normalized counts. The percentage of cells in each population contributing to total *Il18r1* locus expression is provided in the x-axis labels. The dashed line represents equal expression of long and short isoforms; a subset of 5 populations is shown. This is an alternative presentation of the data in **Figure 2D**.

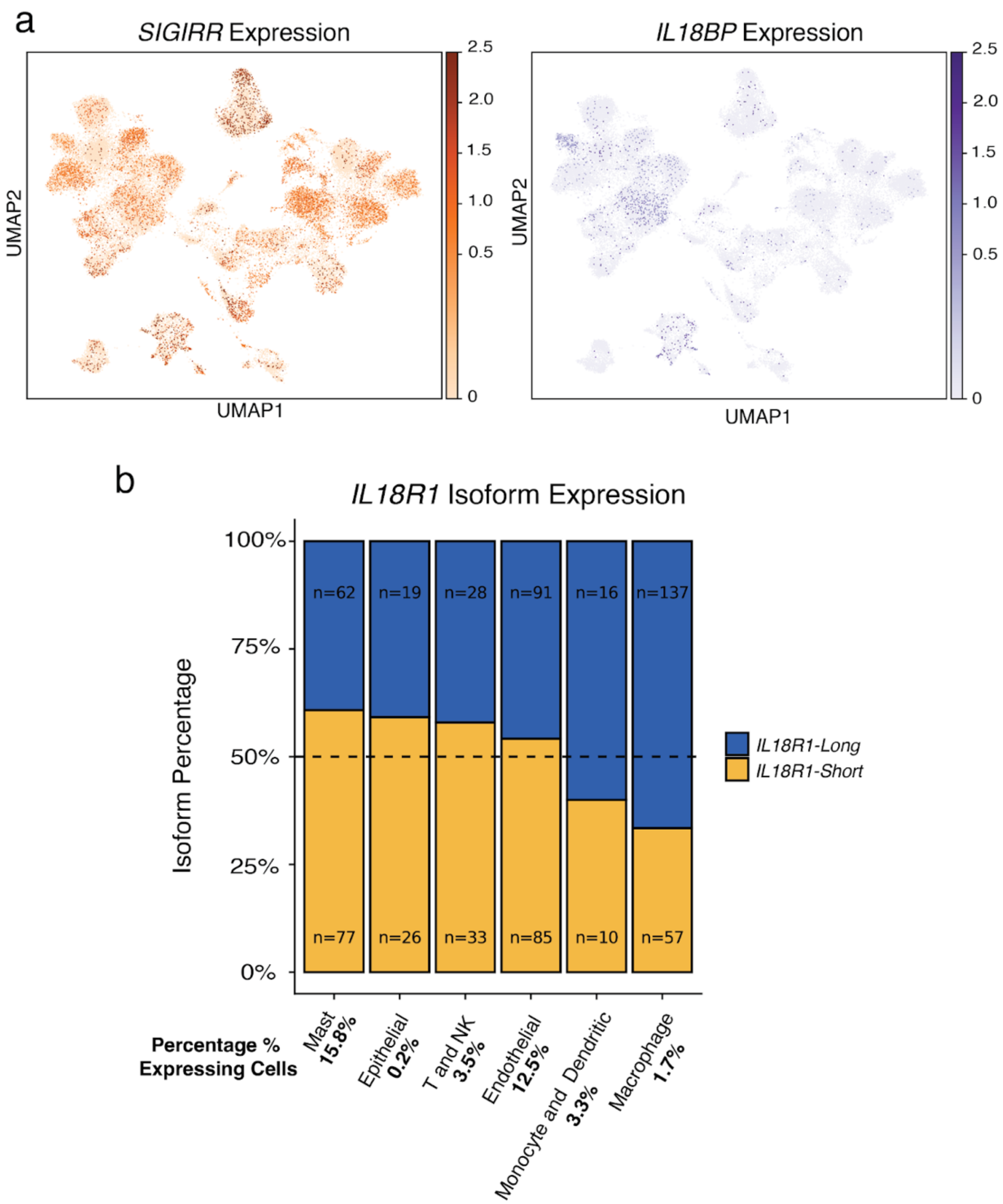


**Fig. S7. Single-cell RNA-seq of human IL18 regulators in the lung. A** UMAP plots showing normalized expression of human *SIGIRR* and *IL18BP* across all profiled lung cells; scale bar represents (log1p (counts per 10K). Cell population UMAP is found in **Figure 2B** and counts for these genes are included in the dot blot in **Figure 2D**. **B** Stacked bar plots showing the average percent expression for *IL18R1* isoforms from normalized counts. The percentage of cells in each population contributing to total *IL18R1* locus expression is provided in the x-axis labels. The dashed line represents equal expression of long and short isoforms; a subset of 6 populations is shown. This is an alternative presentation of the data in **Figure 2D**.

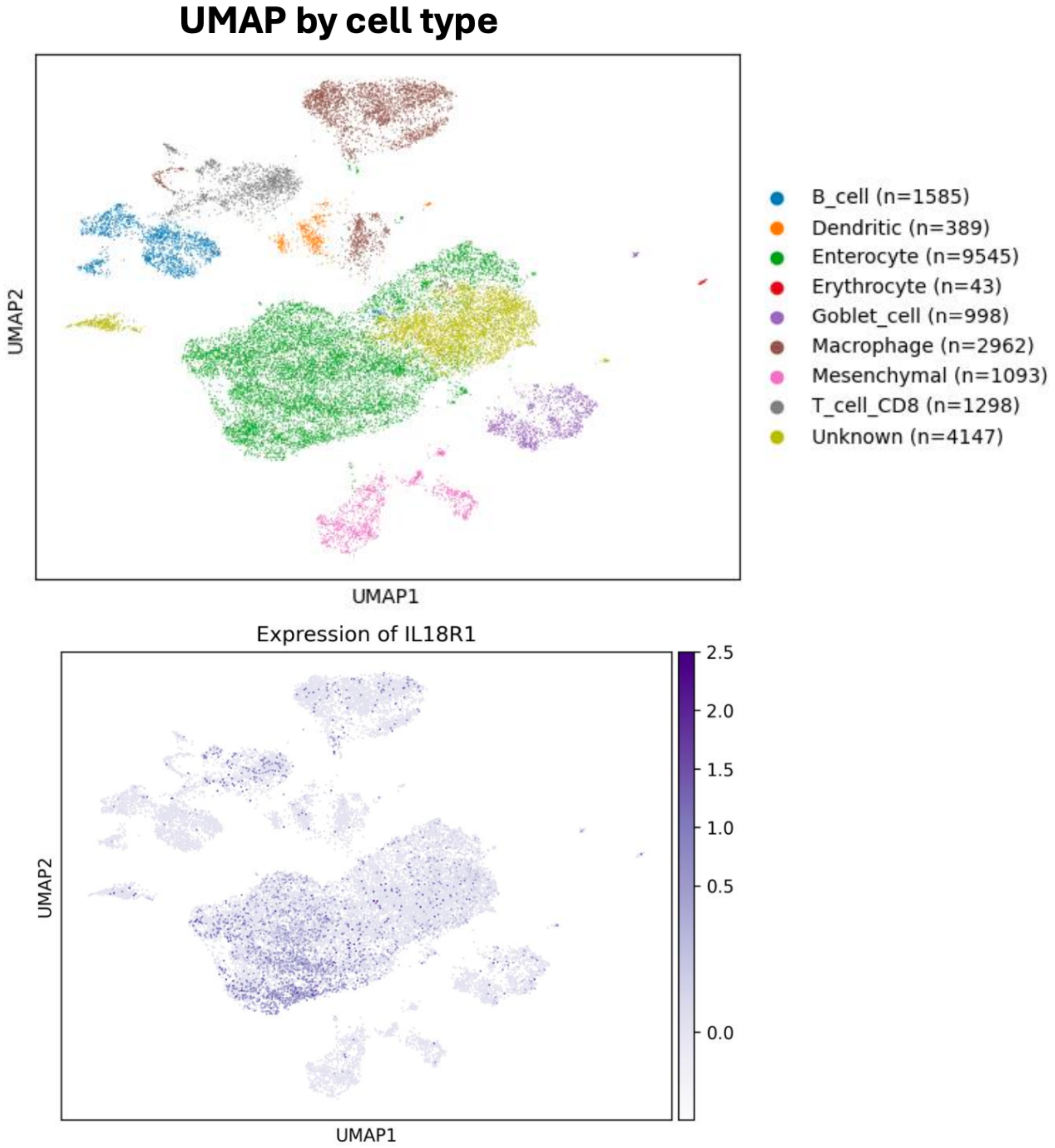


**Fig. S8. Single-cell analysis of *IL18R1* expression in Jamaican fruit bat intestine.** (Top) UMAP plot showing clustered cell populations identified from Jamaican fruit bat intestine single-cell RNA-seq data. (Bottom) normalized expression of *IL18R1*; scale bar represents (log1p (counts per 10K). There was low read coverage at 3'UTRs for long and short isoforms, therefore isoform-level quantification could not be performed. This is supplementary information for **Figure 2.**

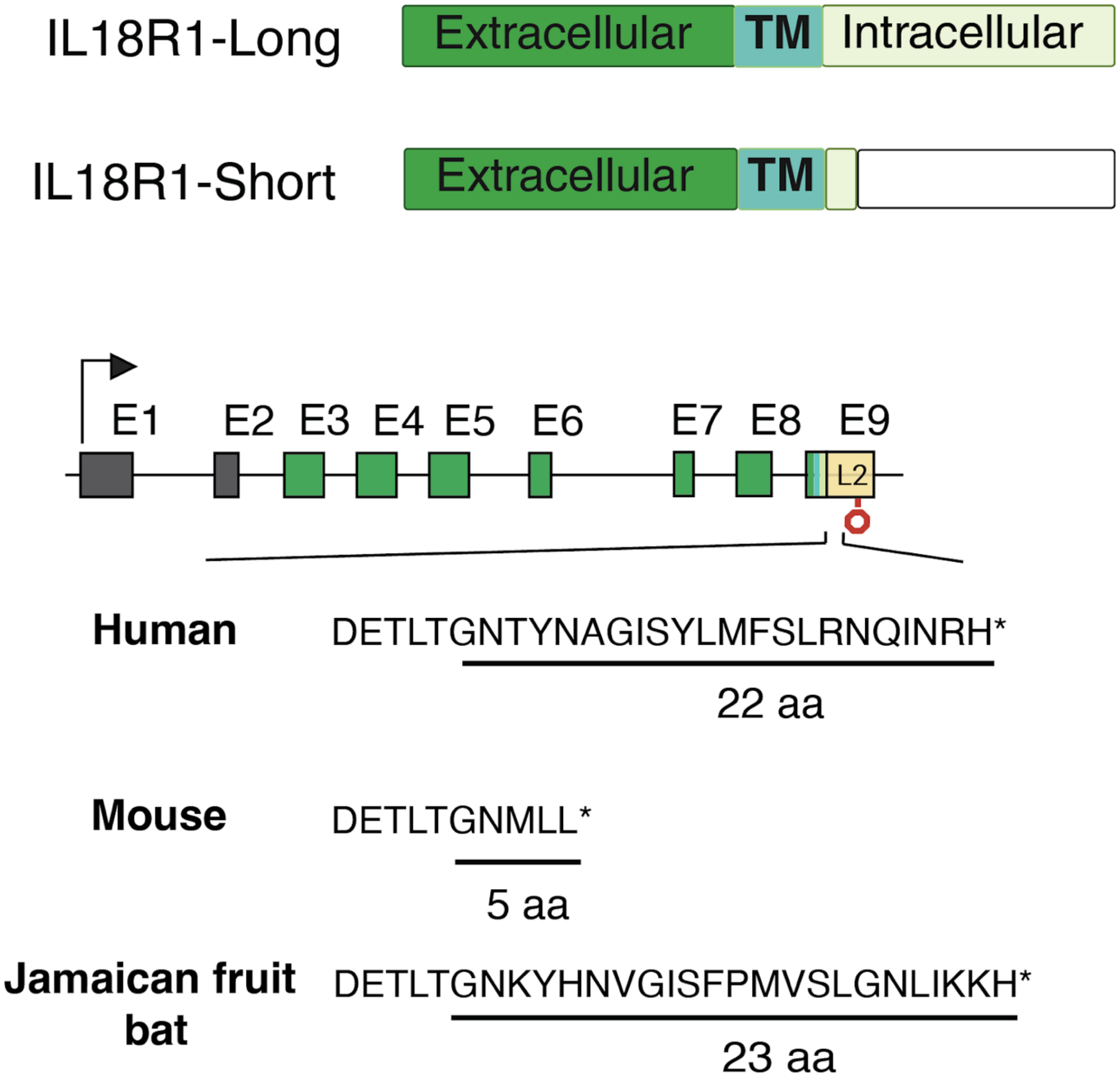


**Fig. S9. IL18R1 truncation across species.** Diagram representations of protein structures for *IL18R1* isoforms. Shown are the unique C-terminal peptide sequences for IL18R1-Short in human, mouse, and Jamaican fruit bat, translated from L2; aa - amino acids. This is supplementary information for **Figure 3.**

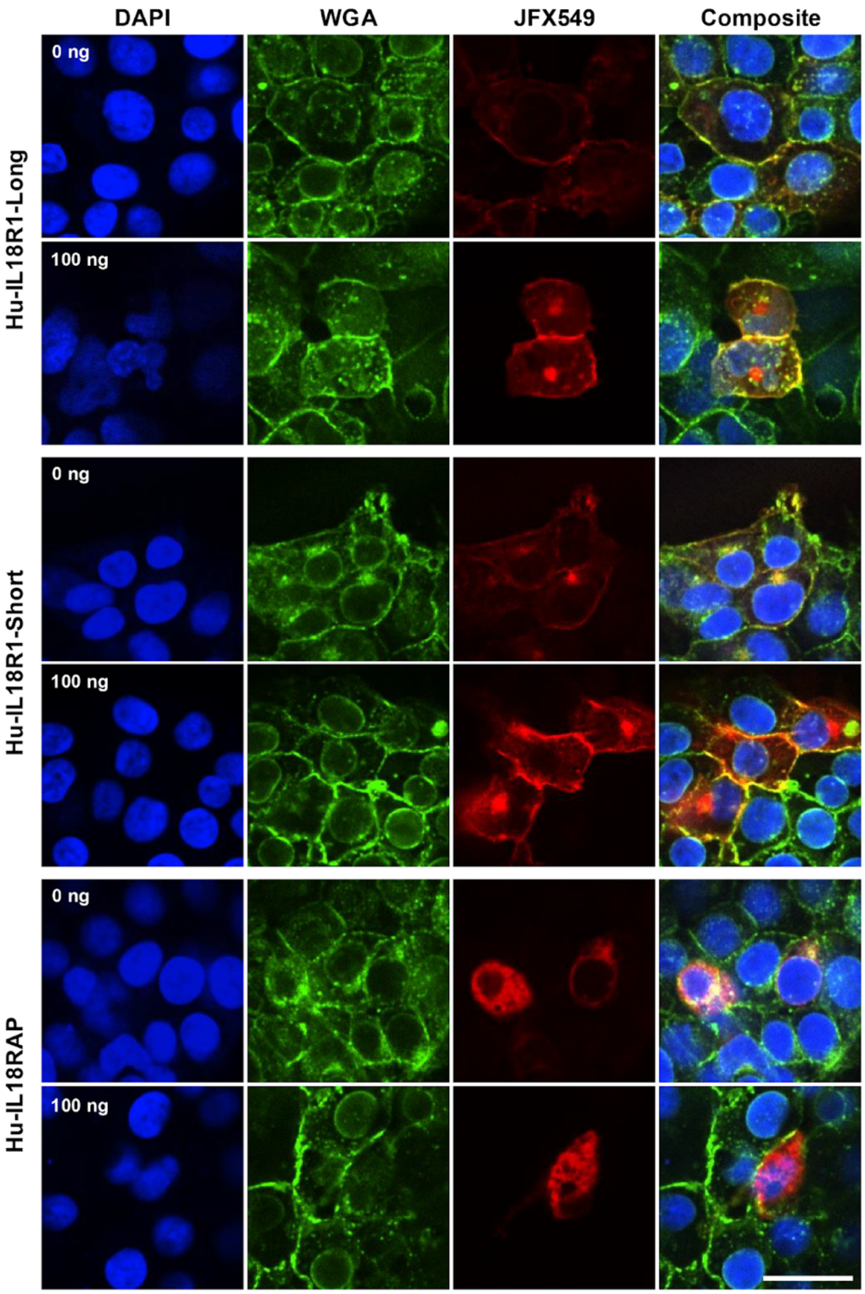


**Fig. S10. Cellular localization of IL18 receptors.** Localization of Human IL18 associated receptors with Human IL18 treatment in HEK293T cells. Representative images of WT, HEK293T cells transfected with either Hu-IL18R1-LongpHTC, Hu-IL18R1-short-pHTC or Hu-IL18RAP-pHTC and treated with 0 ng or 100 ng Hu-IL18 for 30 mins. Images from left to right were taken at 405 nm (DAPI), 488 nm (wheat-germ agglutinin), 561 nm (JaneliaFluor 549, Halotag ligand) and the final image shows all the channels merged. Image brightness for each channel is normalized between treatments for each genotype. Scale bar indicates 50 μm for all images. Supplementary data to **Figure 3A.**

a

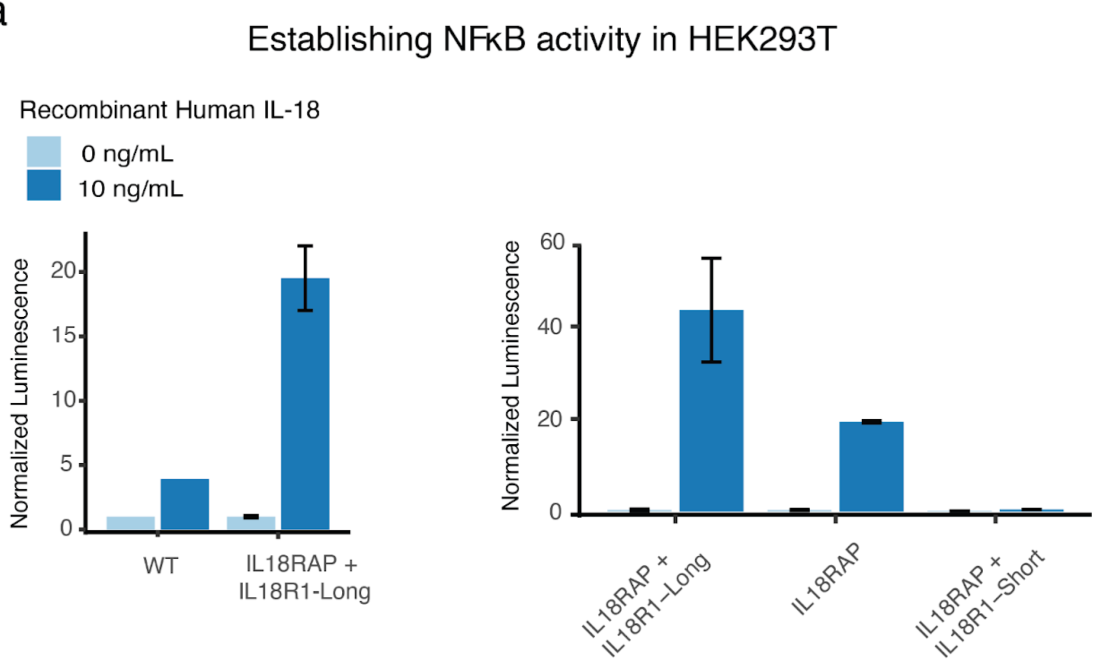


b

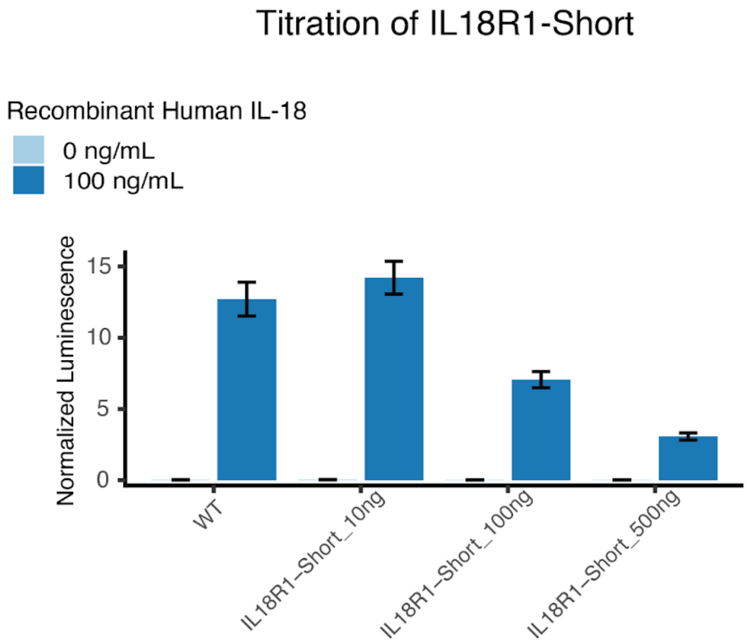


**Fig. S11. Establishing IL18 signaling in HEK cells.** NFκB dual-luciferase reporter assays in human HEK293T in response to human IL-18. Reported values are Nanoluc luminescence normalized to Firefly (NL/FF). **A** Various combinations of human IL18R1-(Long or Short) +IL18Rap; WT = wild type. **B** Titration of human IL18R1-Short. Data is supplementary to the assay in **Figure 3B.** Error bars represent ± standard error (SE) of the mean; N = 4 treatment replicates.

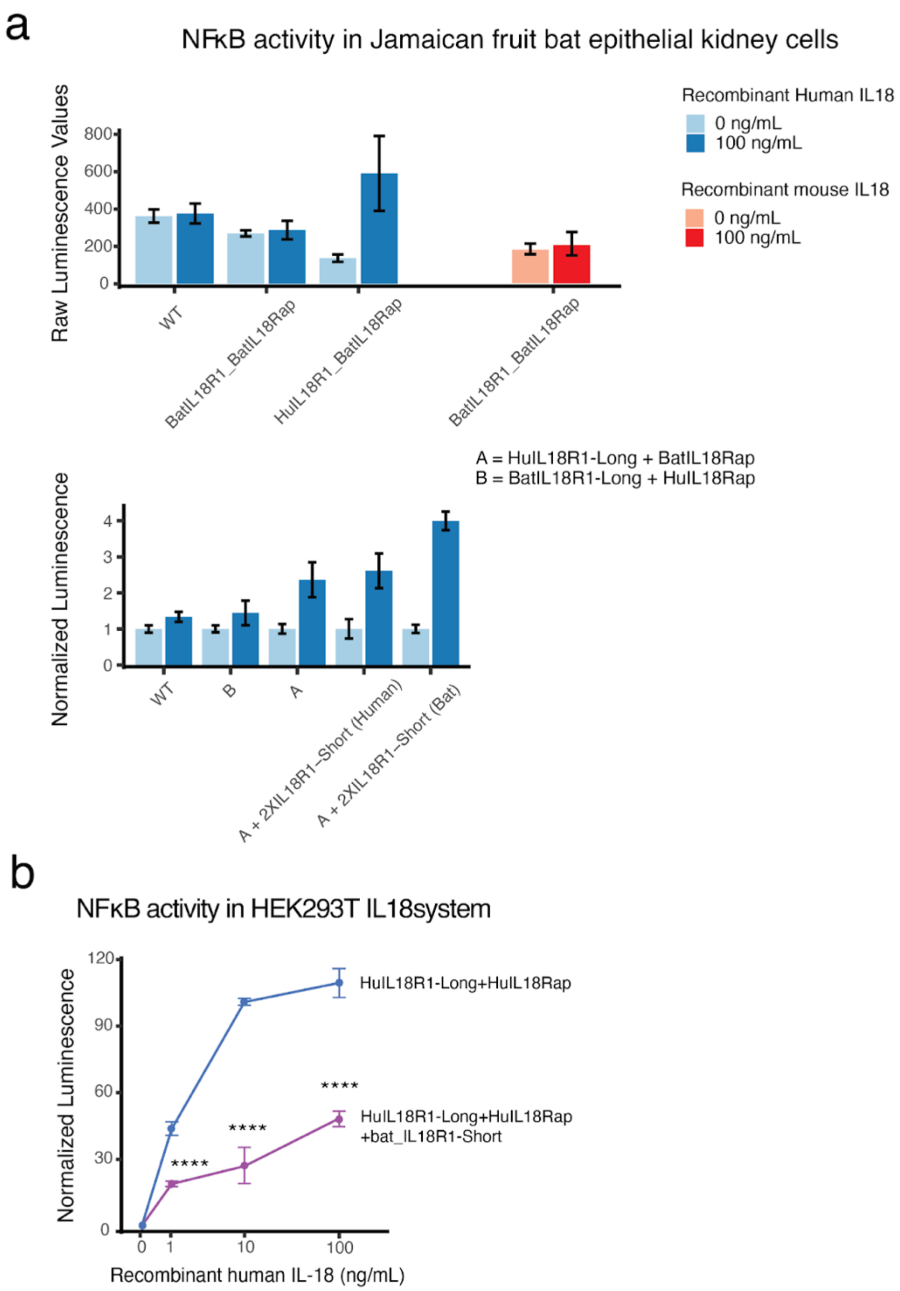


**Fig. S12. Testing bat IL18R1-Short decoy activity. A** NFκB dual-luciferase reporter assays in Jamaican fruit bat epithelial kidney cells in response to human or mouse IL-18; for human IL-18, different combinations of bat and human IL18R1-Long+IL18Rap were tested. Reported values are Nanoluc luminescence normalized to Firefly (NL/FF). Error bars represent ± standard error (SE) of the mean; N = 2 treatment replicates; WT = wild type. **B** NFκB reporter assay of human HEK293T IL-18 reporter system, with or without the over-expression of bat specific IL18R1-Short; data and experimental set-up is from the same assay as **Figure 3B.** P-values were calculated using unpaired two-tailed t-tests (**, p ≤ 0.001; ***, p ≤ 0.001; ; ****, p ≤ 0.0001).

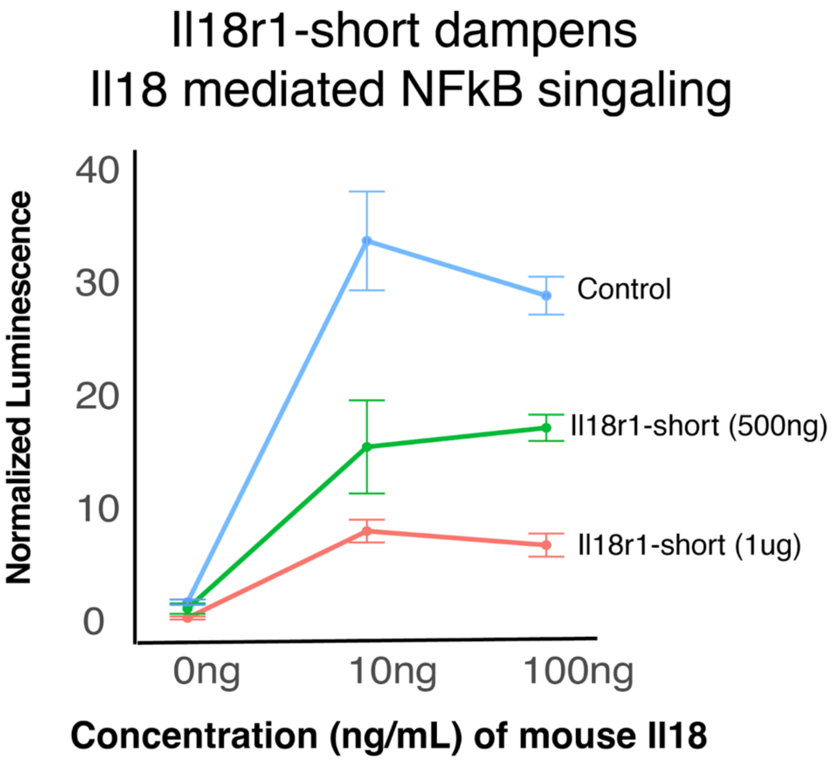


**Fig. S13. Mouse Il18r1-short decoy activity in EL4.** NFκB dual-luciferase reporter assays in mouse EL4 cells in response to varying doses of mouse Il18; varying concentrations of Il18r1-short were tested. NanoLuc luciferase values are normalized to Firefly; NL/FF ratios are normalized to untreated. Error bars represent ± standard error (SE) of the mean; N = 2 treatment replicates. Transfection Control is WT cells transfected with mCherry. Supplementary data for **Figure 3.**

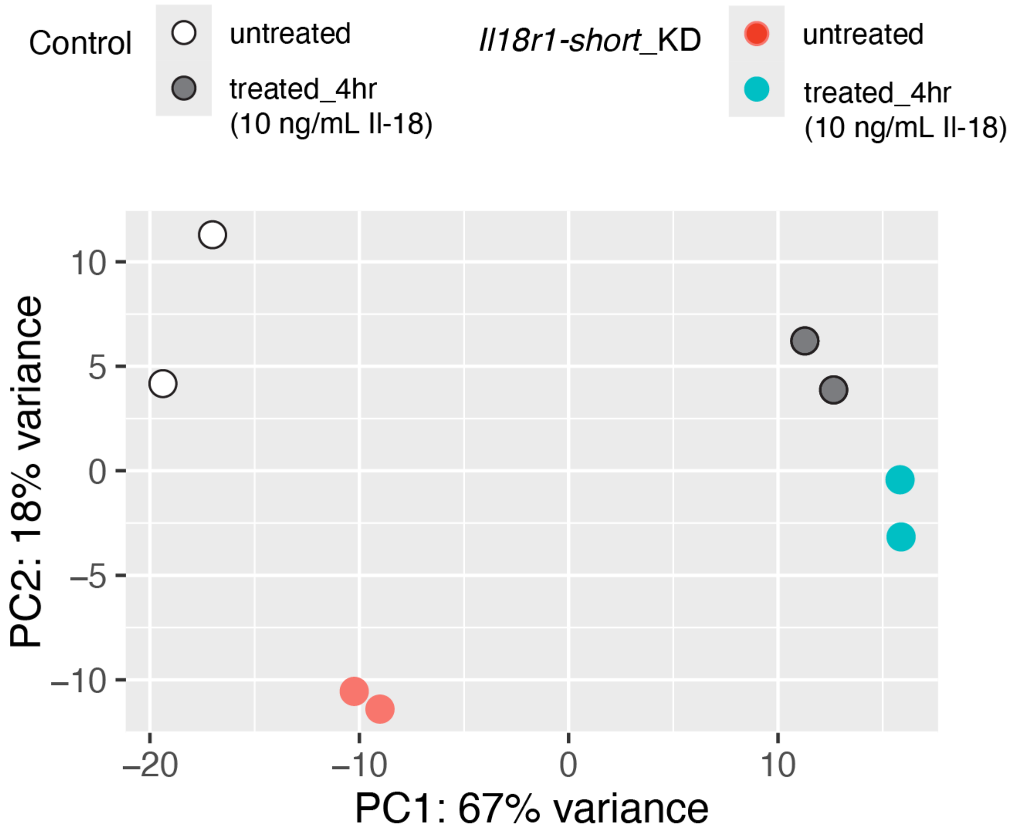


**Fig. S14. Differential gene expression of EL4 Il-18 treated cells.** Principal component analysis (PCA) of transcriptome-wide gene expression in *Il18r1-short* knockdown and control EL4 cells. The data points represent N=2 biological samples for KD and control, untreated and Il-18 treated for 4 hours (10 ng/mL). PCA was performed on variance-stabilized counts (VST) for the 5,000 most variable genes across all samples. Supplementary data to **Figure 4.**

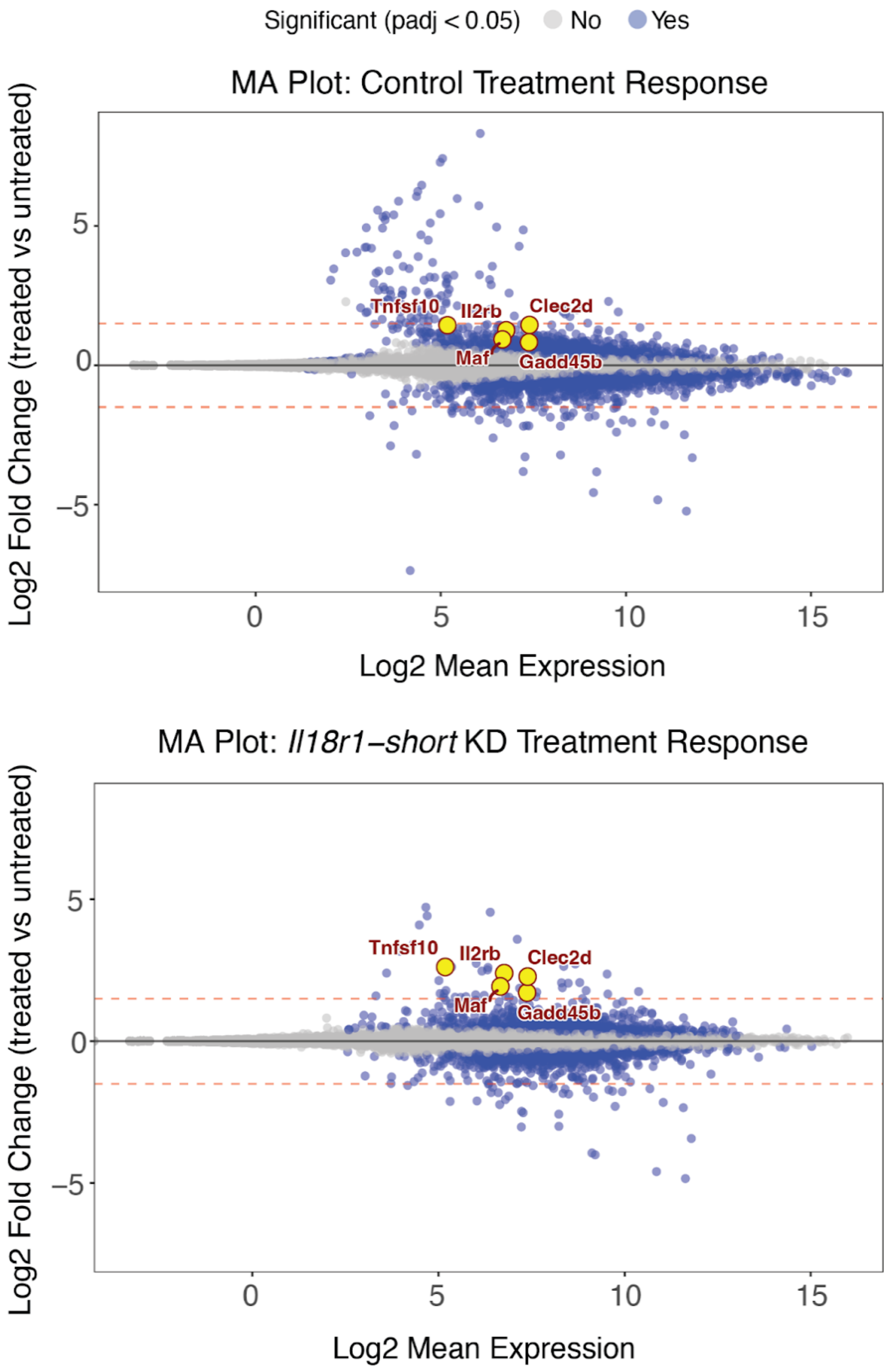


**Fig. S15. Differential treatment response of EL4 cells: *Il18r1-short* KD vs Control.** MA plots of global differential gene expression analysis (DESeq2; design = ~ condition + KD + condition: KD); 4-hour Il-18 treatment (10ng/mL) of mouse EL4 cells with *Il18r1-short* KD or a non-targeting control. The log2fold change of all genes with differential expression are plotted (grey - not significant, blue - significant; p-adj < .05). Red, dashed lines mark 1.5 log2fold difference. Significant genes were further analyzed in **Figure 4**. Genes of interest from **Figure 4C** are labeled here in yellow.

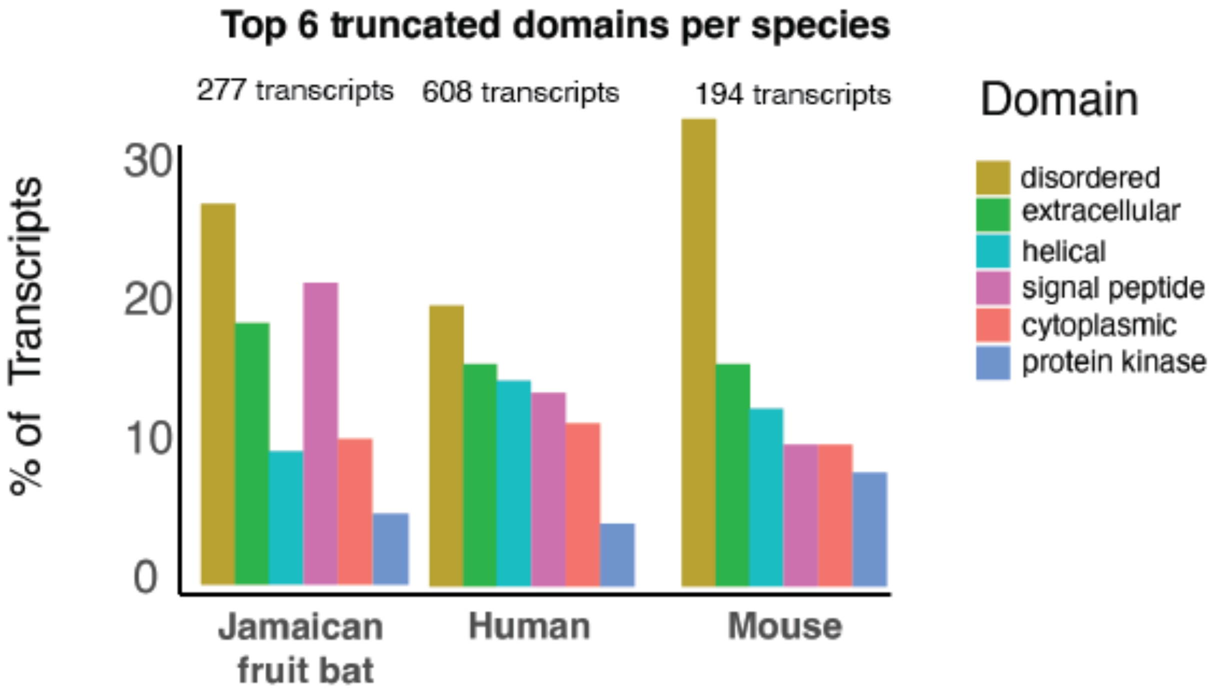


**Fig. S16. Truncated domains in alternative transcripts of immune genes.** Percent of protein domains most deleted in immune genes because of alternative transcription, across species. The top six domain categories were the same across species. The total number of significantly expressed immune gene transcripts (CPM ≥ 5; ≥30% of locus expression) with domain deletions are given at the top. Supplementary data to **Figure 5 A-B.**

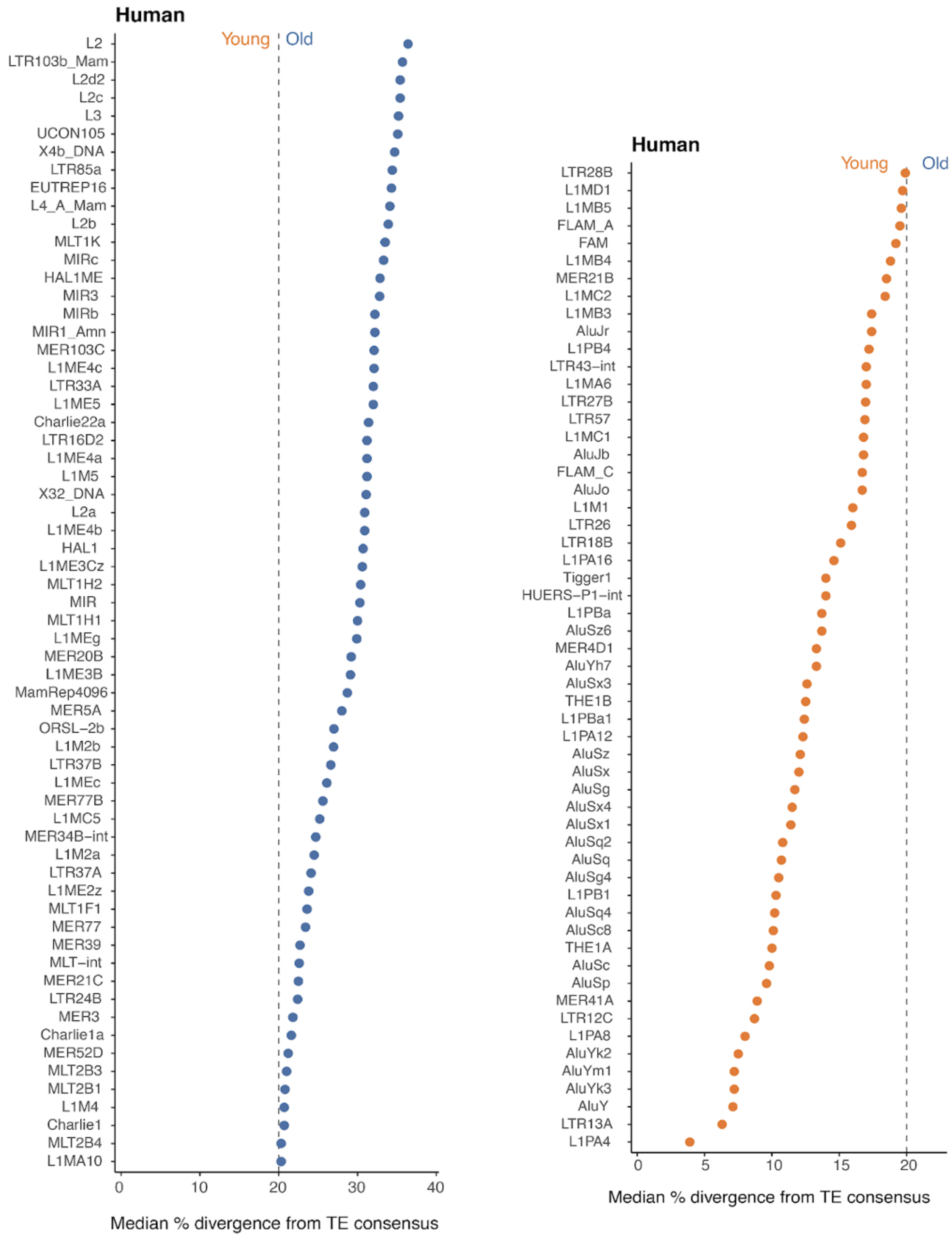


**Fig. S17. Age of human exonized TEs.** Dot plots showing the median percent divergence from consensus sequence for each TE family identified in the human TE-derived alternative transcripts with domain deletion. Divergence values were calculated from all annotated insertions of each TE family in the human genome (hg38), with milliDiv scores divided by 10 to convert to percent divergence. Each dot represents one TE family. The dashed vertical line at 20% divergence demarcates young (orange, <20%) from old (blue, ≥20%) TE families. TE families are ordered by increasing divergence. **Left:** Old TE families (≥20% divergence). **Right:** Young TE families (<20% divergence). Supplementary data to **Figure 5.**

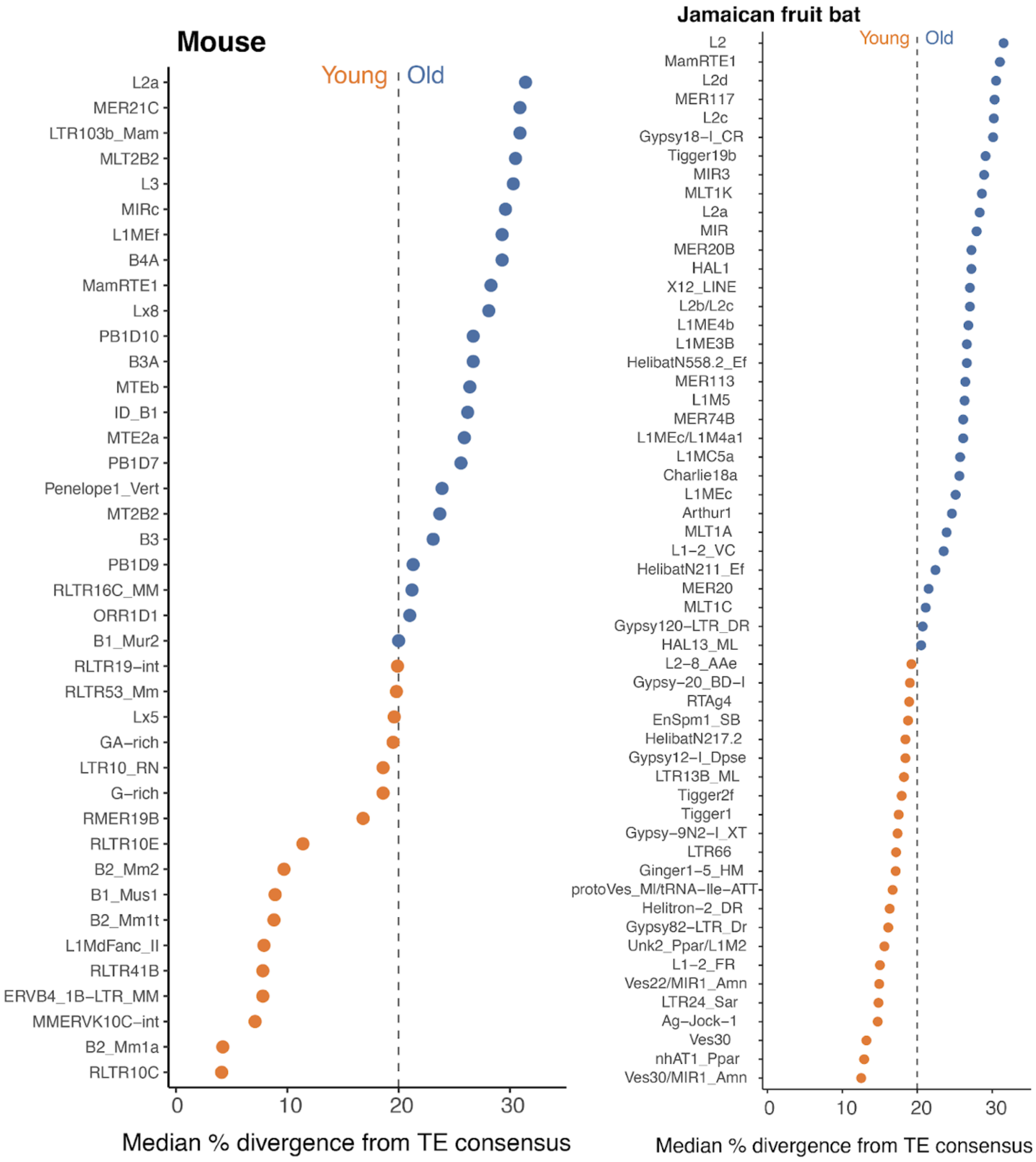


**Fig. S18. Age of bat and mouse exonized TEs.** Dot plots showing the median percent divergence from consensus sequence for each TE family identified in TE-derived alternative transcripts with domain deletion; in the mouse **(Left)** and Jamaican fruit bat **(Right)**. Divergence values were calculated from all annotated copies of each TE family in the respective genome using RepeatMasker output, with milliDiv scores divided by 10 to convert to percent divergence. Each dot represents one TE family. The dashed vertical line at 20% divergence demarcates young (orange, <20%) from old (blue, ≥20%) TE families. TE families are ordered by increasing divergence. Supplementary data to **Figure 5.**

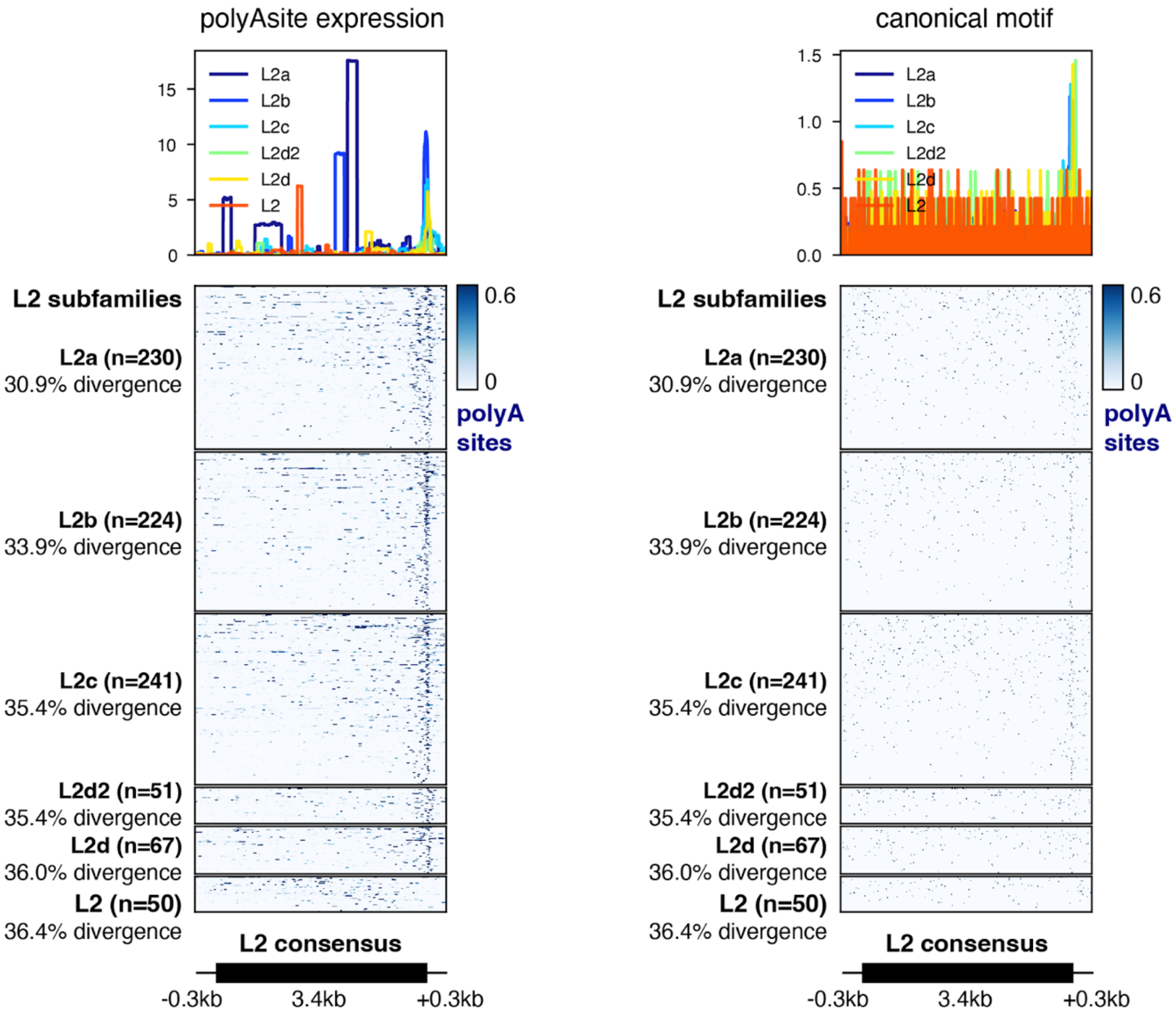


**Fig. S19. L2-PAS motif analysis for human genome.** Comparison of expression-based and sequence-based polyadenylation site enrichment across L2 elements. Left: PolyASite 3.0 single-cell expression signal across L2 loci containing expressed PAS motif (average RPM > 0.9). Right: presence of the canonical AATAAA PAS hexamer across L2 loci, independent of expression data. Both panels show the same L2 elements anchored to the L2 consensus sequence, ordered by increasing sequence divergence. Profile plots above each heatmap show the mean signal across all loci per subfamily. Both approaches independently show enrichment of the PAS motif at the 3' end of L2 elements. Supplementary data to **Figure 5C.**
